# The apoptotic splicing regulators RBM5 and RBM10 are subunits of the U2 snRNP engaged with intron branch sites on chromatin

**DOI:** 10.1101/2023.09.21.558883

**Authors:** Andrey Damianov, Chia-Ho Lin, Jeffrey Huang, Lin Zhou, Yasaman Jami-Alahmadi, James Wohlschlegel, Douglas L. Black

## Abstract

Understanding the mechanisms of pre-mRNA splicing and spliceosome assembly is limited by technical challenges to examining spliceosomes in vivo. Here we report the isolation of RNP complexes derived from precatalytic A or B-like spliceosomes solubilized from the chromatin pellet of lysed nuclei. We found that these complexes contain U2 snRNP proteins and a portion of the U2 snRNA, bound with intronic branch sites prior to the first catalytic step of splicing. Sequencing these pre-mRNA fragments allowed the transcriptome-wide mapping of branch sites with high sensitivity. In addition to known U2 snRNP proteins, these complexes contained the proteins RBM5 and RBM10. RBM5 and RBM10 are alternative splicing regulators that control exons affecting apoptosis and cell proliferation in cancer, but were not previously shown to associate with the U2 snRNP or to play roles in branch site selection. We delineate a common segment of RBM5 and RBM10, separate from their known functional domains, that is required for their interaction with the U2 snRNP. We identify a large set of splicing events regulated by RBM5 and RBM10 and find that they predominantly act as splicing silencers. Disruption of their U2 interaction renders the proteins inactive for repression of many alternative exons. We further find that these proteins assemble on branch sites of nearly all exons across the transcriptome, including those whose splicing is not altered by them. We propose a model where RBM5 and RBM10 act as components of the U2 snRNP complex. From within this complex, they sense structural features of branchpoint recognition to either allow progression to functional spliceosome or rejection of the complex to inhibit splicing.

## INTRODUCTION

Pre-mRNA splicing is a key step in the expression of eukaryotic genes that is catalyzed by the spliceosome, a large multisubunit ribonucleoprotein (RNP) particle. Many essential cellular, developmental, and disease processes are driven by changes in splicing and the expression of alternatively spliced mRNA isoforms. These splicing choices are determined by a large number of regulatory RNA binding proteins that alter spliceosome assembly [reviewed in ^1–4^].

Spliceosome assembly is highly dynamic [reviewed in ^5–7^], and one key early step is the recruitment of the U2 snRNP by factors bound at the 3’ splice site, followed by base pairing of the U2 snRNA with the branchpoint sequence to form the pre-spliceosomal A complex. The U2/pre-mRNA helix contains a bulged branchpoint A residue whose 2’ hydroxyl will be the attacking group in the first catalytic step of splicing. The A complex then progresses through a series of mature spliceosomes with the sequential recruitment and ejection of many additional factors to ultimately catalyze exon ligation.

Spliceosomes must assemble on each intron and are subject to regulation that alters splice site choices according to cellular conditions. Despite much progress in resolving the structures of key spliceosome assembly intermediates and elucidating the mechanisms of catalysis, many aspects of splicing remain poorly understood. Most splicing in the nucleus occurs during transcription on chromatin, conditions that likely differ dramatically from standard in vitro splicing systems. Moreover, the assembly pathway and spliceosome structures have been examined only on a limited number of introns, chosen for their efficient splicing in HeLa nuclear extract. Of particular interest for biology are the many regulatory proteins that alter splicing choices. These factors’ interactions with pre-mRNA’s can be mapped using genomic sequencing approaches, but their interactions with the spliceosome have largely been studied in nuclear extracts and not a native context. New methods are needed for examining regulator/spliceosome interactions in different cellular contexts in vivo.

Two regulators studied both for their biology and the mechanism by which they alter splicing are the proteins RBM5 and RBM10, which are implicated in cancer development, where they alter apoptotic regulation and cell proliferation ^8–12^. RBM10 is also seen to be mutated in the multisystem genetic disorder TARP syndrome ^13,14^. RBM5 and RBM10 share domain structures ^11,15,16^, and both proteins interact with several U2 snRNP-specific or U2-associated proteins ^15,17–20^, and were detected in A- and B-complex spliceosomes ^21^. Despite high sequence similarity of their RNA binding domains, their RNA binding preferences differ ^8,16,22–26^. In vivo crosslinking studies found RBM10 binding to be enriched upstream of branch sites, in proximity to the U2 snRNP ^16,25^. Interestingly, RBM5 crosslinking was more prominent downstream of branch sites and did not consistently map to its target exons. The mechanism by which RBM5 and RBM10 can target particular exons for regulation is not fully clear.

In exploring the interactions of splicing factors within the chromatin compartment of cells, we previously isolated a large, multimeric complex called LASR that binds Rbfox family proteins and contains other splicing regulators ^27^. We were interested in whether other regulatory interactions could be identified in this subcellular compartment, including between splicing regulators and spliceosomal proteins. Examining components of the branch site recognition machinery associated with chromatin, we identified an unusual U2 RNP particle that contains RBM5 and RBM10. We show that isolation of RNA from this particle allows the profiling of intronic branch sites assembled into endogenously formed precatalytic spliceosomes. We examine how U2-bound RBM5 and RBM10 might drive splice site choices through modulation of branchpoint interactions.

## RESULTS

### The splicing regulatory proteins RBM5 and RBM10 are bound within U2 snRNP complexes isolated from chromatin

To assess protein interactions with U2 snRNPs assembling onto nascent RNA, we isolated nuclei, subjected them to gentle lysis, and pelleted the high-molecular weight (HMW) material containing chromatin and nascent RNA. To release spliceosomal components, this pellet was extracted with a cocktail of enzymes digesting RNA and DNA. Examining the sedimentation of spliceosomal proteins and splicing regulators in glycerol gradients identified factors present in higher order molecular complexes. We found that the U2 proteins SF3A3 and SF3B1 were present in a nuclease resistant complex that cosedimented with the splicing regulator RBM5 at approximately 15S. (Figure S1A). These complexes were observed in HMW extract of both mouse brain and human 293Flp-In cells, but not in the soluble nucleoplasmic fraction where the three proteins migrated near the top of the gradient. To characterize these complexes, we generated 293Flp-In cell lines inducibly expressing either SF3A3-Flag or RBM5-Flag under tetracycline control. Induced expression of Flag-tagged SF3A3 reduced the level of the endogenous SF3A3, but did not affect the levels of other SF3A and SF3B proteins (Figure S1B). We then immunopurified the complexes from the peak gradient fractions (Figures S1A and S1C, indicated in blue) on anti-Flag agarose and analyzed their components by gel electrophoresis and mass spectrometry. The protein profiles of the SF3A3-Flag and RBM5-Flag complexes were very similar (Figure 1A and Table S1). These profiles resemble the 17S U2 snRNP ^28^ with a notable lack of the U2A’, U2B” and the Sm core proteins (Figure 1B). Unlike U2 snRNPs characterized from standard nuclear extracts, the U2 in the chromatin fraction of cells has additional components that include the regulatory protein RBM5.

**Figure 1.**
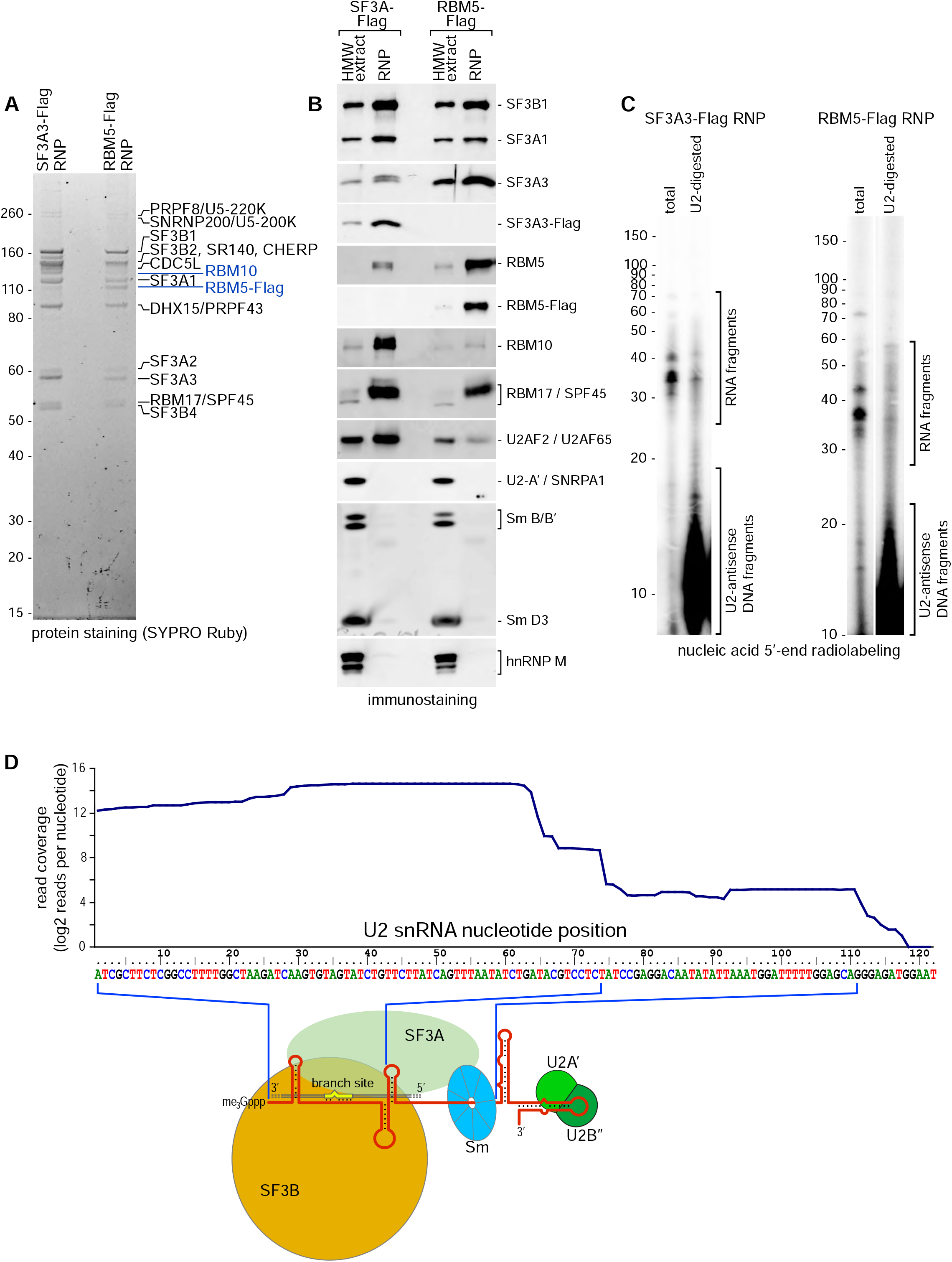
The SF3A3 and RBM5 RNP complexes isolated from HMW extract have similar protein components, and contain U2 snRNA and other RNA fragments. (A and B) Protein profiles of immunopurified SF3A3-Flag or RBM5-Flag-containing complexes from gradient regions indicated in blue in Fig. S1 AC. Proteins are detected either by staining with SyproRuby (A), or immunostaining (B). Lanes marked “HMW extract” in B correspond to 5% input. (C) 5’-radiolabeled RNA, recovered from SF3A3-Flag or RBM5-Flag RNPs, separated by denaturing Urea-PAGE and detected by phosphorimaging. Samples were pretreated with full-length U2 snRNA-antisense DNA and RNase H, followed by TurboDNase digestion (lanes “U2-digested”), or with TurboDNase only (lanes “total”). Detected RNA fragments and short digested DNA oligonucleotides are indicated on the right. (D) U2 snRNA read coverage per nucleotide from isolated RBM5-Flag RNP mapped according to position along the RNA. Only the nucleotides 1-122 are shown since no reads were mapped downstream. The read coverage relative to the overall structure of the 17S and A-complex U2 snRNP in is diagrammed below.

Interestingly, the SF3A3-Flag complex contained limited RBM5 but larger amounts of the related protein RBM10, reflecting the relative abundance of these proteins in the HEK293 cells. Since the RBM5 complex did not contain RBM10, these two paralogous proteins appear to bind U2 in a mutually exclusive manner (Figures 1B). To examine RBM10, we created a cell line expressing RBM10-Flag. In this case, we immunoprecipitated RBM10-Flag from total HMW extract without gradient fractionation (Figure S2A). The profile of proteins coprecipitated with RBM10 looks almost identical to those of the RBM5 and SF3A3 complexes, but with excess RBM10, indicating a mix of free RBM10-Flag and U2 bound protein. The RBM10 also coprecipitated some of the U2 proteins from the nucleoplasm, albeit less efficiently. Thus, the paralogous proteins RBM5 and RBM10 both bind to the U2 snRNP.

The third member of this RBM5 family is RBM6. In a cell line expressing Flag-tagged RBM6, we found the most abundant coprecipitating proteins were the scaffold attachment proteins SAFB and SAFB2 (Figure S3 and Table S1). Despite its sequence similarity and shared domain organization with RBM5 and RBM10, RBM6 did not interact with the U2 snRNP proteins, and appears to have a different function.

Despite extraction with nuclease, all of the isolated U2 complexes retain RNA components. As expected, several discrete bands were identified as U2 snRNA fragments by their sensitivity to RNase H degradation in the presence of U2 antisense oligonucleotide (Figures 1C, S2B). Sequencing RNA from the RBM5 containing U2 RNP, we found U2 snRNA reads were the most numerous among all snRNA reads (Figure S4). We also noted high levels of reads from the U11 and U12 snRNAs (see below). The U2 reads predominantly mapped to the 5’ half of the snRNA, with highest protection from nuclease digestion between U2 nucleotides 27 and 65, which includes the branch site interaction sequence (Figure 1D). The recovered reads drop at the Sm core binding site, and are absent after nucleotide 120. This is consistent with the observed absence of the Sm proteins and the U2A’ and B” proteins that bind in these regions (Figure 1AB).

The U2 Auxiliary Factor U2AF2/U2AF65 was found to interact with RBM5 ^15^, and could potentially recruit RBM5 to the U2 snRNP. However, U2AF2 was sub-stoichiometric in the isolated complexes, and bound to the RBM5-Flag complex at even lower levels than to SF3A3-Flag (Figures 1AB, Table S1). The presence of RBM5 in the U2 complex is thus unlikely to be mediated by U2AF2 binding. Similarly, RBM5 was found to bind the Sm B protein via its OCRE domain ^29^. The Sm core proteins are absent from the U2 complex isolated here and thus are not a contact point for RBM5 (Figure 1B). These interactions may occur in the larger U2/pre-mRNA complexes present prior to nuclease digestion.

Another potential interaction for RBM5 and RBM10 within the isolated U2 RNPs is the RNA helicase DHX15. DHX15 enters the spliceosome as a component of the 17S U2 snRNP and remains associated until after catalysis, when it stimulates disassembly through interaction with the G-patch protein TFIP11 ^28,30^. Several recent studies have also implicated DHX15 and the G-patch protein SUGP1 in a quality control step of early spliceosome assembly ^31–33^. DHX15 was present in the RNPs isolated with Flag-tagged RBM5, RBM10, or SF3A3, but there was limited SUGP1 in these complexes (Figures 1A, S2A). Interestingly, RBM5 and RBM10 also contain G-patch motifs and the RBM5 motif was shown to interact with DHX15 and increase its helicase activity in vitro ^18^. With this in mind, we tested if the G-patch motifs in RBM5 and RBM10 mediated their recruitment to the U2 RNP. We found no differences between the complexes isolated with full length or G-patch deletion mutants of these proteins, indicating that the G-patch was not required for the U2 interaction (Figure S5AB).

To identify residues in RBM5 that mediate interaction with U2, we made a range of additional deletion mutations across the proteins. These mutations removed the known functional motifs within the proteins, including the two RRM domains, the zinc finger, and the OCRE domain. We also deleted the N-terminal 90 amino acids, and the C-terminus that includes the G-patch. All of these deletion mutants maintained their interaction with the U2 complex, copurifying with the same complement of proteins as the full-length RBM5 (Figure 2AB). Finally, we deleted an internal segment of the protein upstream of the C-terminus that contains conserved peptide sequences but was not known to interact with other proteins (Figure S6). In contrast to the other mutations, deletion of RBM5 residues 544 to 703 eliminated its association with the U2 complexes (Figure 2AB). Deletion of the homologous region (residues 655-816) from RBM10 also completely disrupted RBM10 interaction with U2 (Figure 2C). These conserved residues of RBM5 and RBM10 thus include the U2 binding segment of the proteins.

**Figure 2.**
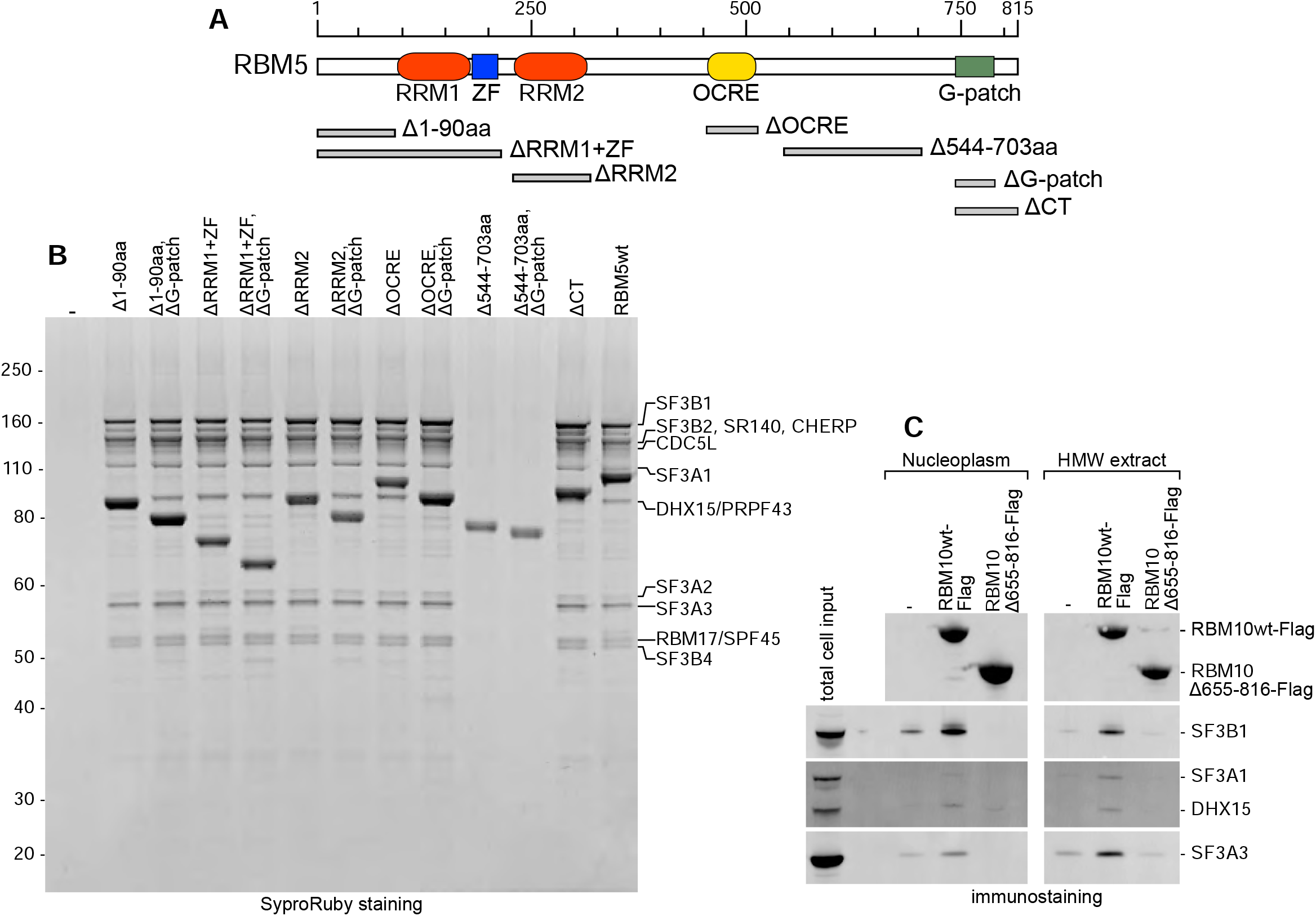
The region between the OCRE and the G-patch motifs in RBM5 and RBM10 is required for interaction with U2. (A) Diagram of RBM5 domains. Significant motifs identified in the CDD/SPARCLE conserved domain database are colored ^68^. Amino acid residue positions are indicated on the top. Grey bars at the bottom indicate sequences deleted in RBM5. Each was deleted independently, or in combination with the G-patch motif. (B) Proteins coprecipitating with the Flag-tagged full-length protein (lane RBM5wt) and with each deletion mutant as indicated on the top. Anti-Flag immunoprecipitations were performed from whole cell HMW extracts. The major coprecipitating proteins detected by protein staining are indicated on the right. (C) Proteins coprecipitating with Flag-tagged full-length RBM10, and RBM10 with amino acids 655-816 deleted (equivalent to RBM5 residues 544-703). Anti-Flag immunoprecipitations were performed from nucleoplasm or HMW extract as indicated, the proteins indicated on the right were detected by immunostaining.

In some sedimentation experiments we noted an additional RBM5 peak, indicating the presence of a smaller-size complex. This peak can be seen in fraction 8 in Fig S1A, but not in Fig S1C. We immunopurified this smaller complex from the gradient fractions. Comparing the protein and RNA composition of this complex to the larger one from fractions 11-13, we found that it lacked the SF3A proteins (Fig. S7A, Table S2) and RNA (Fig. S7B). This smaller complex may be an inconsistent product of RNA degradation and SF3A dissociation from the larger one. The presence of RBM5 in this complex indicates that it likely makes a direct interaction one of the remaining proteins such as the SF3B subunits, but not with the SF3A heterotrimer or the U2 snRNA.

### RBM5 and RBM10 are components of branchpoint-engaged U2 snRNPs across many exons

The U2 complexes isolated from chromatin contained RNA species that remained after U2 snRNA degradation with RNase H (Figures 1C, S2B). To characterize these RNA components, we generated sequencing libraries from the Flag isolated complexes after U2 removal by RNase H. The isolated RNA fragments (28 to 55 nt) were converted to cDNA and sequenced on the Illumina platform. The sample preparations were called RNP-seq if they included a gradient fractionation step, or IP-seq if the RNA was isolated with just a Flag-IP and peptide elution. Mapping the isolated RNA sequences identified them as pre-mRNA fragments. These fragments form tight clusters of mapped reads near the 3’ ends of introns (Figures 3A and S8, see tracks RNP-seq and IP-seq and Table S3 for details). The pre-mRNA sequences overlap branch sites, and thus likely derive from RNA base-paired with U2 snRNA in endogenously formed spliceosomes. These were protected from degradation during extraction from the HMW nuclear material. The peaks of protected branch sites are remarkably homogeneous, with consistent width distribution and few background reads away from branch sites. Virtually all introns in the expressed transcriptome generated a peak of branch site fragments, although the peaks vary in height from intron to intron. Surprisingly, this was seen for all complexes, isolated with SF3A3-Flag, RBM5-Flag, and RBM10-Flag. While some branch site fragments are more abundant in RBM5 or RBM10 complexes, and others more abundant in SF3A3 complexes, we found that virtually all branchpoints were bound by U2 complexes containing RBM5 and RBM10, in addition to SF3A3. These included branch sites for both constitutive and RBM-regulated exons (see below). The reads from the SF3A3, RBM5, and RBM10 complexes together generated 385,806 clusters, 145,934 of which aligned with an annotated branchpoint.

**Figure 3.**
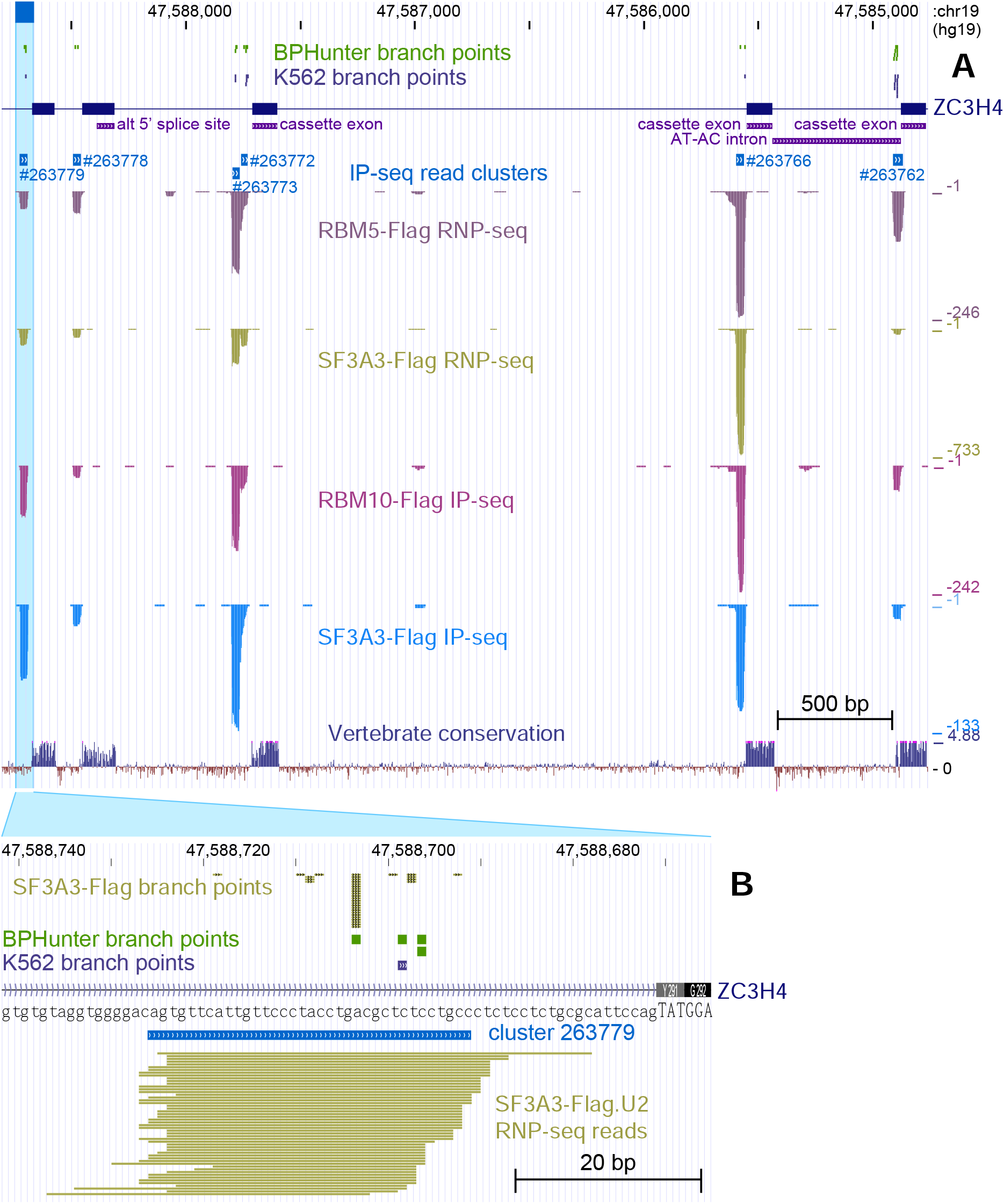
The SF3A3, RBM5, and RBM10 RNPs are bound to pre-mRNA branch sites. (A and B) UCSC Genome browser view of protected RNA sequencing library reads mapped to portions of the ZC3H4 gene. Total read coverage is shown in A below the gene diagram, with individual reads shown in B. Individual tracks show RNA recovered from the SF3A3-Flag and RBM5-Flag RNPs (isolated on gradients, see Figure 1), or RNA coprecipitated with SF3A3-Flag and RBM10-Flag directly from HMW extracts (see Figure S2) (lanes RNP-seq or IP-seq). Significant branch site clusters of overlapping reads are marked in tracks above the peaks. Above the gene diagram are tracks of branch points identified by others ^37,38^. Precatalytic branch points predicted from reads in RNP-seq and IP-seq are shown as vertical bars with height proportional to the frequency of each predicted branchpoint in B (see Figure S10 for details).

The protection of the intronic 3’ ends was not limited to the major-class introns, but also included U12-type introns (shown for a ZC3H4 AT-AC intron in Figure 3A). This was unexpected since the SF3A heterotrimer was found to be absent from the 18S U11/U12 snRNP ^34,35^ and is thought to be replaced with SCNM1 in the minor spliceosome ^36^. Compared to U2 sites, the U12 branch site peaks were more efficiently recovered with RBM5 and RBM10 than SF3A3 (Fig. S9AB). Limited amounts of SF3A3 may be present in minor class spliceosomes at an as yet undefined stage of assembly. RBM10 and particularly RBM5 efficiently recovered branchpoints of minor class introns. It will be interesting to explore whether they play a role in their splicing.

Identified branched nucleotides from available datasets ^37,38^ mapped near the middle of individual reads of the U2-protected RNA fragments (see Figure 3B for example). The reverse transcriptase used to generate the sequencing libraries of these RNAs is expected to terminate at a branched nucleotide. The efficient generation of reads spanning these sites indicates that the RNAs are unbranched, and the purified U2 complexes are derived from precatalytic spliceosomes. Considering the stability of the complexes, with high recovery of branch site fragments over the background, these RNPs most likely resemble A- or B-like spliceosomes, rather than earlier complexes before the U2 snRNP interaction with the pre-mRNA is stabilized.

In A and B precatalytic spliceosomes the pre-mRNA is base-paired to the U2 snRNA with the future branched nucleotide unpaired and bulged from flanking helical stems ^39–43^. To characterize the positions of the future branch nucleotides within our thousands of protected branchpoint regions, we aligned our branch site clusters to datasets of mapped and predicted branchpoints. The mapped branchpoints were identified in K562 and other human cell lines and tissues, but not HEK293 cells ^37,38^. Thus, there were many protected fragments within our data where the branchpoint was not yet known. In a filtered set of clusters each containing a single previously mapped branchpoint (Figure S10A), the branch nucleotide was most frequently located downstream of the read midpoint, within a narrow 50% interquartile range (IQR) (Figure S10B). Focusing on this range and applying a tool for ranking branch site heptamers, we identified the best-fitting putative branch point for each read across all clusters (Figure S10C). We then ranked the most frequently called branchpoints within the reads of each cluster (Figure S10D). These are shown as branchpoint tracks in Figures 3B and S8. The ranks of these predicted branch points strongly correlated with the position of previously mapped sites (Figure S11). The nucleotide frequencies surrounding these predicted branch nucleotides closely matched the previously defined consensus sequences (see below). Thus, we can accurately predict the engaged branchpoints for each intron generating a read cluster in our data.

The sequencing of pre-mRNA fragments bound by U2 snRNPs provides a comprehensive map of branchpoint interactions across the transcriptome. The levels of unspliced introns across a gene are highly variable, with many present at low levels due to their rapid excision ^44,45^. However, the low background of non-branchpoint reads and the precise alignment of the protected fragments allows identification of the U2 binding sites for nearly all the introns in the expressed RNA. The tracks presented here derive from one replicate of three for each condition. Together these replicates generated datasets of 24-84 million reads per experiment, over 65% of which mapped to branch site clusters and provided sensitive detection of the branchpoints (Table S3). The branchpoint peaks vary in height for each intron across a gene transcript. Some of this variation presumably results from different intron excision rates leading to branchpoints spending greater or lesser time within pre-catalytic spliceosomes. However, this does not appear to be the only factor affecting the peak heights, and we have not identified simple rules that determine the recovery of each branchpoint. Overall, this profiling approach allows comprehensive assessment of U2 assembly and accurate branchpoint prediction across the transcriptome.

### Identification of RBM5 and RBM10 regulated splicing events

To examine the effects of RBM5 and RBM10 on individual exons across the transcriptome, we established a system to identify RBM5 and/or RBM10-regulated splicing events. We engineered HEK293 cells with the endogenous genes disrupted and replaced with either with RBM5-Flag or RBM10-Flag, expressed from a tetracycline-inducible promoter. Using CRISPR-Cas9 to disrupt both alleles of the two genes, we could isolate cells lacking RBM5, but were unable to disrupt the reading frame of both RBM10 alleles in this RBM5^-/-^ background. We isolated a clone with both RBM5 alleles and one RBM10 allele inactivated, and with one RBM10 allele expressing limited amounts of a mutant RBM10 mRNA containing a cryptic exon (CE) that restores the reading frame after the targeted deletion (Figure S12). These RBM5 null, RBM10 mutant cells (RBM5^-/-^; RBM10^-/CE^) exhibited markedly slower growth than wildtype cells, and this growth was rescued by expression of either RBM5 or RBM10. We then compared gene expression and splicing in the mutant cells with cells rescued by either protein.

We isolated polyA+ RNA and sequenced it from four replicas each of the parental mutant cells and the RBM5-Flag and RBM10-Flag rescued cells induced to express the proteins at similar levels (Figure 4A, see also Figure S13). The RNAseq data were analyzed by rMATS to examine RBM dependent splicing ^46^. These analyses identified many RBM5 and RBM10-dependent alternative splicing events, including cassette exons, alternative 5’ and 3’ splice sites and retained introns (Figure S14, Tables S4, S5). Cassette exons were the most numerous. Previous studies identified smaller sets of RBM5 and RBM10 dependent exons in HeLa and mouse ES cells ^16,25^. There was notable overlap between the different studies of RBM10 regulated exons. Of the exons affected by RBM10 depletion in HeLa, 23% were also RBM10 regulated in our HEK293 system, including a well characterized RBM10 target exon in Numb. We found that RBM5 and RBM10 predominantly downregulated their target exons and that a substantial fraction of these targets was coregulated by both proteins (Figure 4B).

**Figure 4.**
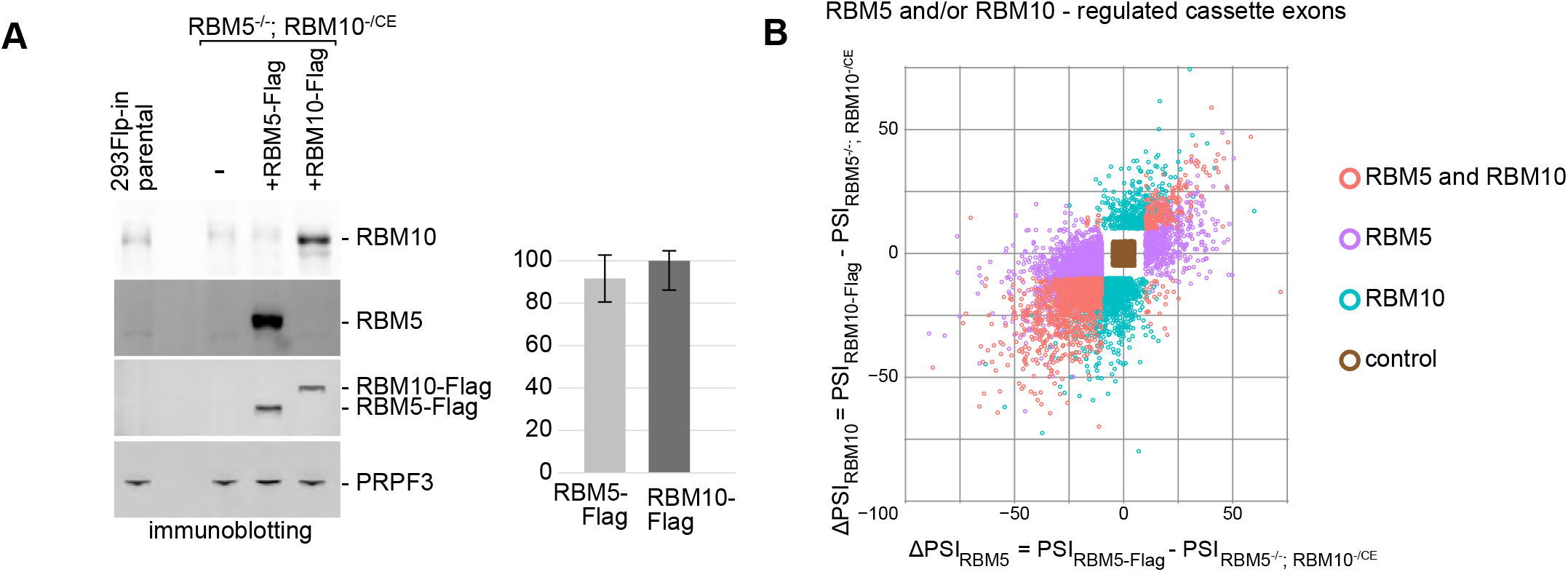
RBM5 and RBM10 repress splicing of many cassette exons. (A) RBM5 and RBM10 expression in 293Flp-In parental cells, in the RBM5^-/-^; RBM10^-/CE^ cell line, grown in presence of 0.6 ng/mL Doxycycline for 72 hours, and in RBM5^-/-^; RBM10^-/CE^ cells transfected with RBM5-Flag or RBM10-Flag transgenes induced for the same time with 0.6 ng/mL or 0.06 ng/mL Doxycycline, respectively. Whole cell RIPA lysates were subjected to immunoblotting. RBM5 and RBM10 were detected with antibodies recognizing endogenous epitopes and with Flag antibody as indicated. PRPF3 was also detected as an internal control. The immunoblot for one of four replicas is shown at the top, with the mean normalized expression of the Flag-tagged proteins from all replicas is graphed at the bottom (error-bars indicate standard deviation). (B) Comparison of RBM5 and RBM10 splicing activities. Cassette exons identified by rMATS ^46^ as altered by RBM5 and/or RBM10 expression compared to the RBM5^-/-^; RBM10^-/CE^ cells are graphed by scatter plot. X axis reports ΔPSI values for RBM5 relative to the parental cells; Y axis reports the RBM10 ΔPSI values. Cassette exons with |ΔPSI| ≥ 10 and FDR ≤ 0.05 are color-coded to indicate regulation by RBM5 and/or RBM10. A control set of cassette exons in the central box with |RBM5 ΔPSI| < 5 and |RBM10 ΔPSI| < 5 is shown in brown.

Comparing branch site clusters isolated from RBM5-Flag or RBM10-Flag complexes with those isolated from SF3A3-Flag complexes, we found that the peak heights upstream of cassette exons did not correlate with the effects of RBM5 or RBM10 on splicing (Figure S15). We also did not find notable differences in the specific branch nucleotides selected by these complexes. The highest-ranking predicted branch point nucleotides matched at 60-65% of the RBM5, RBM10, or SF3A3 clusters. Neither this match frequency nor changes in predicted branch site position correlated with RBM5 and/or RBM10 downregulation (Figure S16).

We next looked for motifs enriched in sequences adjacent to the branch sites of regulated exons. In A- and B-like spliceosomes nucleotides adjacent to the U2-paired sequence are contacted by SF3B and SF3A proteins ^47–49^. Regulatory motifs could also be present at sites more distal to the protected nucleotides of the branch site. The upstream and downstream regions differ in nucleotide composition due to the downstream polypyrimidine tract, the intron terminal AG dinucleotide, and exonic sequences. To separately examine these regions, we aligned the intronic sequences at the highest-ranking branchpoint predicted for RBM5-Flag or RBM10-Flag clusters, and then defined intervals upstream and downstream (see diagram in Figure S17A). Since the distance from the branchpoint to the exon can vary, we limited the downstream intronic region to range from 10 to 46nt. This excluded a small fraction of branchpoints at greater or lesser distance from the downstream exon. We separately analyzed the sequences at the intron-exon boundaries, including the last five intronic and the first twenty exonic nucleotides. For each set of branchpoint-adjacent sequences, we used STREME to compare the motif frequencies within the RBM5 and/or RBM10 regulated targets to those of a control set of cassette exons unaffected by RBM5 or RBM10 ^50^. We found that both RBM5- and RBM10-downregulated targets are enriched for a variety of C-rich motifs upstream and downstream of the branchpoint. These enrichments were more significant for the RBM5 regulated exons. However, these C-rich sequences have only limited similarities to previously identified RBM5 binding motifs ^22–24^. There was also enrichment for a G-rich motif in the region 16-36 nt upstream that is similar to an RBM5 binding motif identified by RNA Bind-N-Seq, although this enrichment did not have as strong a p-value (Figure S17A).

To test the contribution of individual sequence motifs in regulating RBM5- and RBM10-dependent exons we constructed several minigenes carrying target exons, and confirmed their regulation by RBM5 and/or RBM10 in the 293Flip-In cells. We then mutated individual sequence motifs in these minigenes to assess their role in splicing regulation. Overall, we mutated 8 sequence elements in 3 different RBM5/10 target minigenes. Mutations in the C- or G-rich motifs upstream of the branchpoint altered the overall inclusion of most target exons but did not alter their response to RBM5/10 (Figure S17B). Similarly, disrupting C-rich motifs downstream of the branch site of some target exons stimulated their baseline level of splicing, but did not affect RBM5/10 regulation (Figure S17B); One exception was an exon in DPP7 where mutation of the downstream C-rich motif (mt1) resulted in nearly 100% baseline inclusion that was minimally repressed by RBM5 (Figure S17B). Mutation of combinations of elements had no greater effect than single mutations. Moreover, many RBM5 and RBM10-responsive exons identified in the RNA-seq analysis did not have these motifs. Such motifs may affect the efficiency of spliceosome assembly without providing specific contacts for RBM5 and/or RBM10. Instead, RBM5 or RBM10 may alter spliceosome activity by direct interaction with the U2 snRNP. Interestingly, previous analyses of an exon in FAS indicated that RBM5 was altering its splicing without a direct interaction with the pre-mRNA.

### Splicing regulation by RBM5 and RBM10 requires interaction with U2

To test how the loss of U2 binding affects the splicing regulatory activities of RBM5 and RBM10, we attempted to express proteins defective for U2 interaction in RBM5^-/-^; RBM10^-/CE^ cells. However, unlike what was seen for the wildtype proteins, the cells harboring these RBM5 or RBM10 mutant transgenes exhibited a severe growth defect. The limited number of recovered clones had very slow growth rate and were difficult to maintain. This indicated that the U2 binding mutants had a severe loss of function but precluded assaying their splicing activity in this system. As an alternative assay, we compared the activities of full-length and mutant proteins in the parental 293Flp-In cells, which continue to express wildtype RBM5 and RBM10 from the endogenous genes (Figure 5A). Assaying splicing of target exons by RT-PCR at multiple protein concentrations, the RBM5 deletion mutants had greatly reduced activity on most target exons compared to the full-length proteins, and in many cases were completely inactive (Figures 5B and S18). The changes in activity in the RBM10 deletion mutants were less consistent, possibly due to higher endogenous RBM10 levels in these cells. For example, Exon 6 of the TRPT1 gene was repressed by RBM10Δ655-816, but not by RBM5Δ544-703 (Figure 5B). For NUMB exon 9, which was moderately repressed by wildtype RBM10 but not RBM5 in this system, both RBM5 and RBM10 deletion mutants had the opposite effect and promoted exon 9 splicing (Figure 5B). These varying responses of some exons indicate the RBM5 and RBM10 proteins may act by multiple mechanisms.

**Figure 5.**
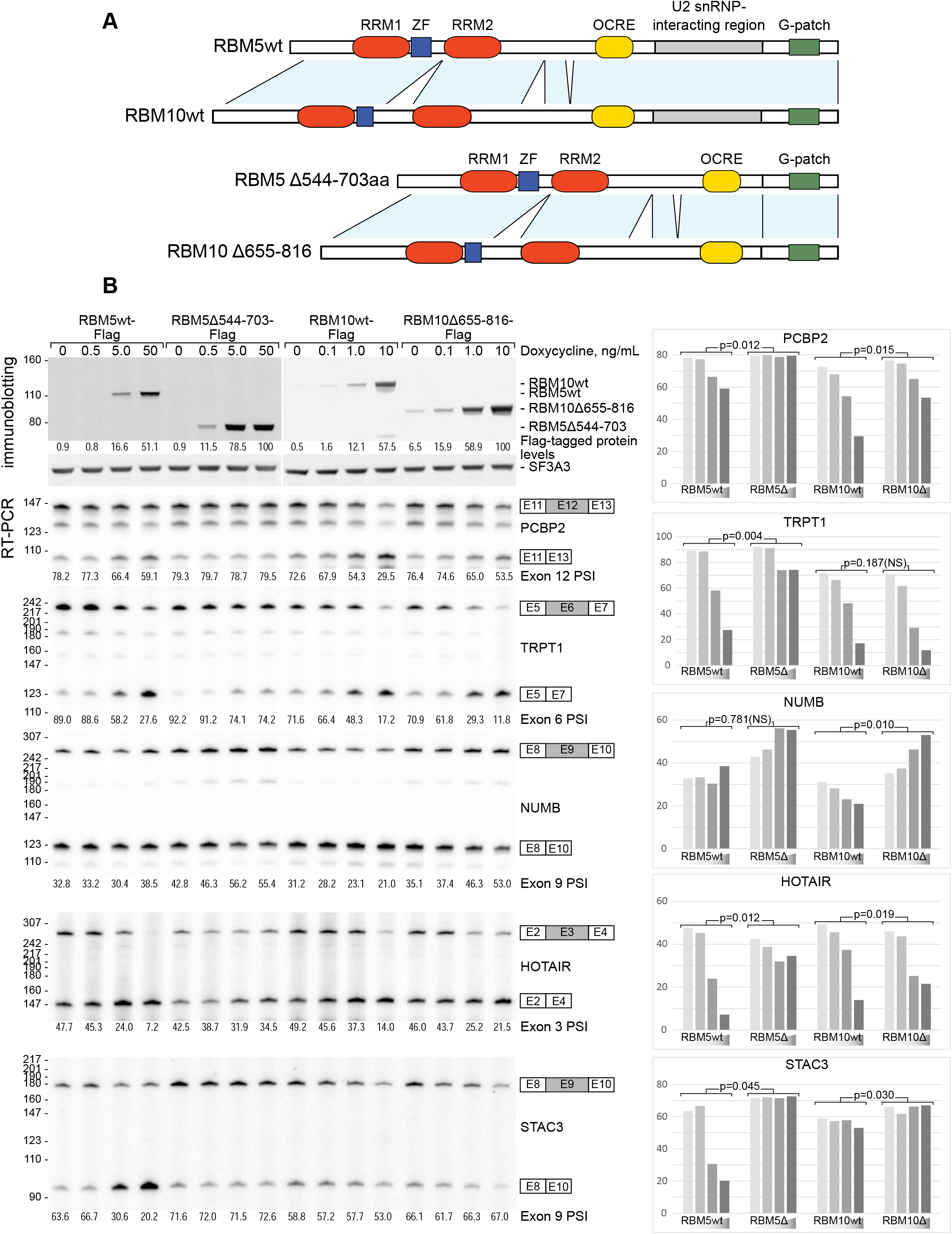
RBM5 and RBM10 require interaction with U2 snRNP to regulate splicing. (A) Diagram indicating known motifs in RBM5 and RBM10, and the peptide segment required for interaction with U2 snRNP. (B) 293Flp-In cells were induced to express Flag-tagged RBM5, RBM5 Δ544-703, RBM10, or RBM10 Δ655-816 as shown in A. Expression of each Flag-tagged protein was quantified after immunostaining, with the relative expression levels indicated below. SF3A3 expression served as a normalization control. Splicing of endogenous PCBP2 exon 12, TRPT1 exon 6, and NUMB exon 9, HOTAIR exon 3, and STAC3 exon 9 were assayed 48 hour post-induction by RT-PCR using radiolabeled flanking primers followed by denaturing PAGE and phosphorimaging. The quantified exon inclusion levels are indicated below, and graphed at the right. P-values indicate the significance of the splicing activity differences between the full-length and deletion mutant proteins on each target. These activities were expressed as regression slope for the exon inclusion as a function of Flag-tagged protein expression. NS indicates not significant.

Overall, the results indicate that a portion of the RBM5 protein located between the OCRE domain and the G-patch is required both for its interaction with branch site-bound U2 snRNPs and for its splicing repressive activity on most exon targets.

## DISCUSSION

### In vivo isolation of early spliceosome complexes

We identified a U2 RNP particle that can be extracted from the chromatin fraction of cells using nuclease but without denaturants. This complex contains multiple U2 snRNP proteins, including SF3A and SF3B, and the 5’ portion of the U2 snRNA apparently base-paired to intron branchpoints. Unlike U2 snRNP’s and early spliceosome complexes isolated from standard nuclear extracts, the particle also contains the regulatory proteins RBM5 and RBM10. Within this U2 snRNP particle, both the branch site and the U2 snRNA are protected from RNase digestion, indicating it derives from an early spliceosome where the base-pairing of the branch site has been established. The presence of the SF3A and SF3B complexes, which are removed by DHX16 during formation of the B* complex, is consistent with the particle being an early assembly intermediate. The lack of branched nucleotides in the protected intron RNAs also indicate the particles were isolated prior to the first transesterification reaction ^51,52^. From these considerations, these U2 complexes likely correspond to spliceosomes in the A, B, or B^act^ stages. They could also derive from an assembly intermediate not yet described in vitro.

### Profiling U2/intron interactions in vivo

The RNA fragments protected within the isolated U2 complexes precisely map to intron branch sites across all of the expressed transcriptome. The cluster signal intensities largely correlate with gene expression levels, but vary among branch sites from the same transcripts. This variation did not limit the sensitivity of branch site identification. At the sequencing depth of our analysis, A/B^act^-like spliceosomes were detectible from even weakly expressed transcripts showing exon RPKM of less than 0.1 in standard polyA+ RNAseq data. Using this approach, we unambiguously identified over 140,000 branch site clusters located upstream of annotated exons.

We find that the future branch nucleotide or nucleotides of each intron can be accurately predicted from the position of branch site motifs within the U2-protected pre-mRNA fragments. Since they cover nearly all the expressed introns in the cell, these predicted sites are expected to be more comprehensive than those where direct sequencing is used to detect intron lariats ^37,38,53^. These predicted sites agree well with the physically mapped branched nucleotides in available datasets, but could likely be further improved using more sophisticated branchpoint-calling approaches that recognize alternative U2-branch site pairing modes ^53^.

In addition to annotated branchpoints, we also detect a large number of clusters with lower signals that are located deeper within introns. Public sequence data indicate that some of these are branch sites of poorly spliced introns. These unannotated sites of U2 binding can allow detection of low abundance mRNA isoforms, or of stalled precatalytic complexes that don’t result in intron removal. Interestingly, we also detect about 7,000 clusters overlapping with 5’ splice sites. These protected 5’ splice sites could derive from the same precatalytic spliceosomes as the branchpoints but which survive RNAse treatment with lower yield. In agreement with this, the isolated material contains substoichiometric amounts of other snRNAs as well as U5, U4, and U1-specific proteins. At the current sequencing depth, the numbers of 5’ splice site reads and clusters is limited, but future biochemical optimization may allow more comprehensive profiling of 5’ sites, using approaches similar to that described here. More efficient mapping of these 5’ splice sites may reveal if the pre-mRNA is base-paired with U1 or U6, and thus more precisely identify the stage of the isolated spliceosomes. In earlier work using this isolation strategy, we identified interactions between splicing regulatory proteins in the chromatin fraction of the nucleus that had not been observed in nuclear extracts ^27^. In the future, we will be excited to test additional tags and extraction conditions in applying this method to studies of nuclear regulatory processes.

### Splicing regulation by RBM5 and RBM10

We identify the splicing regulators RBM5 and RBM10 as components bound at high stoichiometry to the U2 snRNP/branchpoint complex. Splicing regulators have largely been studied through their interactions with the pre-mRNA. These pre-mRNA binding events are thought to direct changes in splice site choice through altering early steps in spliceosome assembly. In contrast, we find that RBM5 and RBM10 incorporate into spliceosome complexes on the nearly complete set of branch sites without an apparent dependence on interactions with snRNA or pre-mRNA. This could allow them to alter splicing as U2 snRNP-associated proteins late in spliceosome assembly. Their interactions within the branch site complex could allow them to sense conformations that favor or disfavor further spliceosome assembly on particular introns.

Introns with multiple branchpoint locations were observed in the U2-protected read clusters. In these cases, individual branch points were often preferentially isolated with RBM5 or RBM10 compared to SF3A3. This is consistent with the idea that RBM5 and/or RBM10 binding is sensitive to the particular conformation of the bound U2 snRNP. On the other hand, we did not observe changes in alternative branchpoints on exons subject to RBM5 and RBM10 regulation. Thus, these proteins repress splicing at a later stage spliceosome than the isolated RNPs. It will be interesting to apply this approach to testing the roles of other U2-associated proteins in branch point selection, as well as to examining other patterns of splicing such as alternative 3’ splice sites.

We find conserved segments of RBM5 and RBM10 that interact with U2 are required for cell growth and for their ability to regulate cassette exons. We identified a large set of cassette exons regulated by both RBM5 and RBM10, as well as exons sensitive to one protein or the other. A more limited overlap between RBM5 and RBM10 targets was observed in a previous study ^16^. This is likely due to differences in the experimental systems: detection by microarray rather than RNAseq, and knockdown of proteins by RNAi rather than CRISPR-Cas9 gene deletion and transgene reexpression. RBM10 was absent from U2 snRNPs isolated with tagged RBM5 and vice versa, while both proteins are present but substoichiometric in the complexes isolated with tagged SF3A3. These observations are consistent with the two regulators being mutually exclusive in their interaction with U2. This, along with the overlap of their target exon sets, may indicate a shared mechanism on at least a subset of their targets. It will be very interesting to narrow down the residues of RBM5 and RBM10 involved in this interaction, and to identify their binding partner(s) on U2, which may be one of the SF3B proteins.

The G-patch motifs of RBM5 and RBM10 suggest possible mechanisms of repression. G-patch motifs act as essential coactivators of DEAH-box helicases ^54,55^. DHX15 in the isolated U2 RNP is proposed along with the G-patch protein SUGP1 to mediate a quality control step of early spliceosome assembly ^31–33,56^. The RBM5 G-patch could play a similar role in activating DHX15 to abort spliceosome assembly on certain exons before the first catalytic step. This hypothesis is compatible with studies indicating that RBM5 inhibits splicing by preventing the association of the tri-snRNP ^15^. The isolated U2 complexes also contain two other G-patch proteins, CHERP and RBM17/SPF45. CHERP and RBM17 have been shown interact with the DHX15, with each other, and to co-regulate alternative splicing ^17,57,58^. All these proteins offer many possibilities for regulatory steps in both the quality control of early spliceosome assembly on constitutive introns, and in the selective rejection of particular complexes to control the choice of splice sites and exons.

We identified enriched RNA sequence motifs adjacent to the branchpoints of RBM5 and RBM10 repressed exons, but experiments to mutate these elements did not strongly support their function as binding motifs for the proteins. This is consistent with the regulation of FAS exon 6 by RBM5, where a direct pre-mRNA binding site was also seen to be lacking. Interestingly, introns with C-rich polypyrimidine tracts were found to splice more slowly than those with U-rich tracts ^44^. C-rich polypyrimidine tracts have also been identified as features in the splicing targets of DDX39B, another RNA helicase associated with U2 ^59^. Thus, it is possible that RBM5 inhibits splicing of its targets by disrupting spliceosomes that are slow to transition from A to B* or C complexes.

Taking all the results together, we favor a model where RBM5 and RBM10 act primarily as U2-associated splicing repressors. These proteins are components of the branch site recognition machinery in A or B-like prespliceosomes assembled on nearly all exons. RBM5 or RBM10 could potentially disrupt this complex by activating the DHX15 helicase or otherwise preventing productive spliceosome assembly on particular exons. The specificity of their targeting could be influenced by branch site-proximal sequences that affect assembly kinetics rather than being directly recognized by the RNA binding domains, although direct pre-mRNA interactions are not ruled out. As integral parts of the assembling spliceosome, these factors differ from other well studied splicing regulators and it will be interesting to examine the interactions of RBM5 and RBM10 within the U2 branchpoint complex in more detail.

### Significance and limitations of this study

We describe an experimental approach that allows some of the first biochemical characterizations of endogenously formed spliceosomes. We demonstrate the versatility and sensitivity of this method both for understanding protein regulators of splicing and for the transcriptome-wide mapping of assembling spliceosomes on nascent RNA. We show that two splicing regulators, RBM5 and RBM10 are integral spliceosome components that repress splicing primarily through interactions with the branch site-recognition machinery.

Currently this approach is limited to cells engineered to express epitope-tagged proteins, and we are working to develop antibodies that allow the isolation of untagged RNP complexes. We are also optimizing the extraction method to broaden its application to additional spliceosomal components and to other informational processes in the cell nucleus.

## PUBLIC DATA SUBMISSION

The IP-seq, RNP-seq, and RNA-seq data is available at NCBI GEO under the accession number GSE240608.

## ACKNOWLEDGMENTS

We thank Manuel Ares Jr. (UCSC) and Timothy W. Nilsen (Case Western University) for comments on the manuscript. This work was supported by NIH grants R01GM127473 (A.D.), R35GM136426 (DLB), and R21HG012624 (DLB).

## STAR METHODS

### Gene expression vectors

RBM5, RBM6, and RBM10 cDNAs were obtained by RT-PCR with 5’ and 3’ UTR primers from total HEK293 RNA. These were sequenced and confirmed to encode proteins identical to GeneBank sequences NM_005778 RBM6, NM_001167582, and NM_001204467, respectively. Human SF3A3 was obtained from plasmid DNA carrying an insertion identical with NM_006802. The protein coding regions were fused with C-terminal FLAG tag sequence and inserted in pcDNA5/FRT/TO vector (Thermo Fisher Scientific). DPP7 splicing reporter minigene containing the full-length sequences of exons 5,6, and 7 and the intervening introns was obtained by PCR from HEK293 genomic DNA. This was inserted into the NheI and ApaI sites of pcDNA3.1 vector (Thermo Fisher Scientific). Sequences and descriptions of mutant derivatives are listed in Extended Experimental Procedures.

### Cell culture

Stable HEK293 lines expressing RBM5, RBM6, RBM10, and SF3A3 proteins were prepared using the Flp-In™ T-REx™ System (Thermo Fisher Scientific). An RBM5/10-deficient clone (RBM5^-/-^; RBM10^-/CE^) derived from this cell line was obtained by pairs of CRISPR/Cas9-guided deletions of RBM5 exons 8 and 9 and RBM10 exon 5 using pX330 vector ^60^. Targeted sequences were as follows: RBM5 (TGCTCCCTGCCCCAGTTAGT_AGG; TAAGTTCATACACGATCTTT_TGG), RBM10 (TCGCTGGCAGCCACGAAGTA_AGG; GCTGAGGCTGGCCACTTGAA_GGG). Transfection of minigenes was performed in 40%-confluent monolayer cell cultures with Lipofectamine 2000 (Thermo Fisher Scientific), 10% minigene-encoding pcDNA3.1 vector, and 90% empty vector for 4 hours. Cells were then harvested 48 hours post-induction with doxycycline, samples for RNA and protein analysis were collected.

### RT-PCR

Total RNA was extracted with TRIzol (Thermo Fisher Scientific) from pelleted cells. DNA was removed with TURBO DNase (Thermo Fisher Scientific). Reverse transcription was carried with SuperScript IV (Thermo Fisher Scientific) and d(`T`)_20_ reverse primer for endogenous mRNA, or with the pcDNA3.1-specific reverse primer GCAACTAGAAGGCACAGTCG for RNA transcribed from minigenes. cDNAs were amplified for 15-20 PCR cycles with Phusion Hot Start II DNA Polymerase (Thermo Fisher Scientific) and 5’-radiolabeled primers, resolved by denaturing PAGE, and detected by phosphorimaging in Amersham Typhoon and quantified in ImageQuant (GE Healthcare Bio-Sciences). Primer and minigene sequences are listed in Extended Experimental Procedures.

### RNP sample preparation

Nucleoplasm and HMW material from highly purified nuclei were obtained as described ^27^ in lysis buffer supplemented with 0.5 mM CaCl_2_. Extraction of nuclease-resistant complexes was performed as before, in presence of approximately 5 U/μl of Benzonase Nuclease (Millipore-Sigma), 0.04 U/μl TURBO DNase (Thermo Fisher Scientific), 0.01 U/μl RNase A, and 0.4 U/μl RNase T1 (RNase Cocktail, Thermo Fisher Scientific). Whole cell soluble and HMW fractions were obtained similarly from pelleted cells lysed for 5 min in lysis buffer, and subjected to RNA and DNA degradation as above. Complexes from extracted material were separated in 10-30% glycerol gradients containing 20 mM HEPES-KOH pH 7.5, 150 mM NaCl, 1.5 mM MgCl_2_, 0.5 mM DTT, and 1x Complete Protease Inhibitor Cocktail (Millipore-Sigma) by centrifugation in Optima XL centrifuge and SW41Ti rotor (Beckman Coulter) for 17h at 32,000 RPM, 4°C. Gradients were manually fractionated top to bottom into 0.5 mL aliquots. Fractions of interest were dialyzed against buffer lacking glycerol and protease inhibitors twice for 30 min at 4°C in Slide-A-Lyzer MINI Dialysis Devices, 10K MWCO Membrane (Thermo Fisher Scientific).

Extracts from nuclear or whole cell fractions, and dialyzed gradient fractions were incubated overnight at 4 °C with 5-7.5 μl packed-volume M2 FLAG agarose beads (Millipore-Sigma), washed four times with wash buffer (20 mM HEPES-KOH pH 7.5, 150 mM NaCl, 1.5 mM MgCl_2_, and 0.05% Triton-X100), and eluted for two hours at 4°C in presence of 150 ng/μl of 3xFLAG peptide (Millipore-Sigma).

### Protein analysis

Immunopurified proteins were separated by SDS-PAGE and detected by immunoblotting with fluorescently-labeled secondary antibodies (GE Healthcare Bio-Sciences), or stained with SYPRO Ruby (Thermo Fisher Scientific). Images were visualized in Amersham Typhoon and quantified as above. Primary antibodies are listed in Extended Experimental Procedures. Proteins from these samples were also pelleted with four volumes of acetone at -20°C overnight, washed with 85% Ethanol, and subjected to Liquid Chromatography Tandem Mass Spectrometry (LC-MS/MS). Spectral counts from proteomic analysis of the indicated purifications are shown in Tables S1 and S2.

### RNA analysis

RNA from deproteinized samples, each in a triplicate, was allowed to hybridize with U2-antisense DNA oligonucleotide (reverse-complimentary to GeneBank sequence NR_002716) and incubated with 0.375 U/μl RNase H (New England BioLabs) for 20 min at 37 °C. DNA was digested with TURBO DNase simultaneously with removal of 5’ and 3’ phosphates with FastAP alkaline phosphatase (Thermo Fisher Scientific) for 45 min at 37 °C. Analytical aliquots (5-10%) of RNA were 5’-radiolabeled by incubation in presence of γ-[^32^P] ATP (PerkinElmer) and T4 Polynucleotide kinase (New England BioLabs), resolved by denaturing PAGE, and imaged in Amersham Typhoon. Sequencing libraries were obtained from preparative aliquots (90-95%) of RNA using iCLIP reagents ^61^ and similar strategy with changes to allow for reactions in solution instead of on carrier surface. Detailed protocol is described in Extended Experimental Procedures. These libraries were sequenced on a HiSeq4000 (Illumina). PCR duplicates were removed using random barcodes. Unique reads from RBM5 library prepared without selective U2 snRNA degradation mapping to snRNA sequences were identified with BLAST ^62^, allowing for one mismatch per 20 nt. Unique reads from all libraries were mapped to human genome hg19/NCBI37 using Bowtie2, excluding partially aligned reads and allowing for one mismatch per 20 nt ^63^. Regions of at least ten overlapping reads in each experiment were defined by YODEL ^64^, and trimmed to exclude terminal sequence with coverage lower than 50% of the maximum. Overlapping regions from one or more RNP-seq or IP-seq experiment were then combinded to define clusters. These were further categorized as 5’ splice site or branch site-associated if being the closets to a 5’ or a 3’ end of an annotated intron, with maximum allowed distance of 10 and 100nt, respectively. Cluster RPKM was calculated with SeqMonk software (v1.45.4, Babraham Institute).

The distribution of known branched nucleotide positions on reads was determined from a subset of clusters, unambiguously identified as branch site-associated, with width of less than 53 nt. Clusters overlapping with less than one, or with multiple branch point identified in ^37,38^ were removed. Also excluded were clusters where read coverage at the branch point nucleotide was less than 85% of the maximum coverage for the cluster. Median branch point positions and IQR intervals as fraction of read length were determined for each library. Using these distributions, a putative branch point was then assigned to each read from clusters other than categorized as 5’ splice site-associated. These were chosen as the position within the IQR for the read with the highest HSF branch site score ^65,66^. Finally, the putative reads within each cluster were ranked by the fraction of supporting reads out of all cluster reads. For branch site-associated clusters on U12-type introns ^67^, a single U12-type branch point with the highest similarity to the U12 branch site consensus sequence out of all predicted branch points was called separately.

### Genome-wide splicing analysis

Total TRIzol-extracted RNA was treated with TURBO DNase. cDNA libraries were prepared from polyA+ RNA with TruSeq Kits (Illumina). Alternative splicing was analyzed by rMATS ^46^ and expressed as changes in percent-spliced-in values (ΔPSI). Exons showing splicing change (|ΔPSI|>10 with FDR less than 0.1) between control RBM5^-/-^; RBM10^-/CE^ cells, and cells expressing RBM5-Flag or RBM10-Flag were considered regulatory targets.

### Motif enrichment analyses

Motifs enriched upstream or within RBM5- and/or RBM10-downregulated cassette exons were determined using STREME ^50^. A set of cassette exons not regulated by RBM5 or 10 (|ΔPSI| < 5 and FDR > 0.05) served as control. Intronic regions were selected based on the position of rank1 branch point from the corresponding branch site cluster.

## SUPPLEMENTARY FIGURE LEGENDS

**Figure S1.**
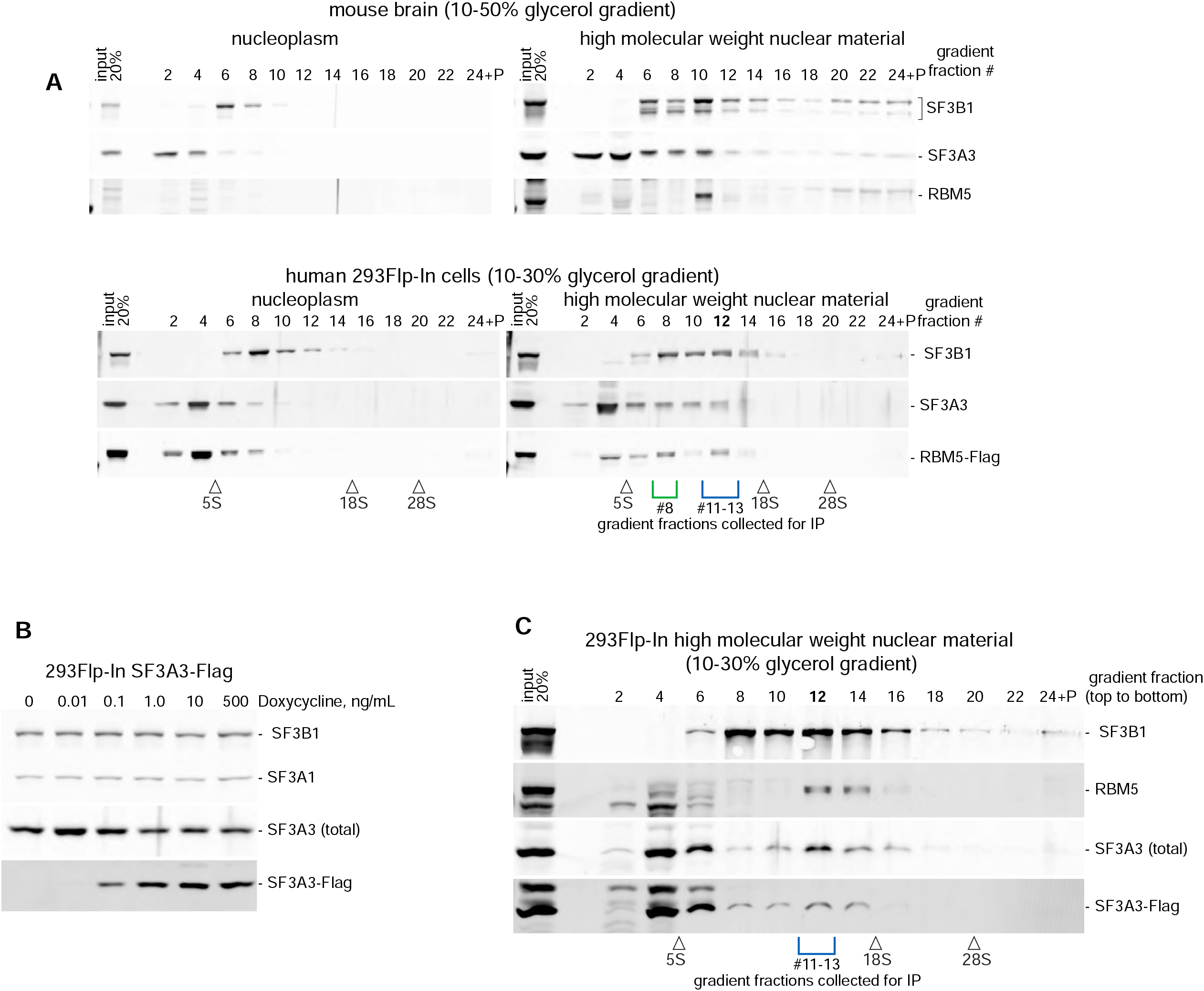
Cosedimentation of RBM5 with U2 snRNP proteins in chromatin extracts (Related to Figure 1) (A) The soluble nuclear fraction and HMW extract from mouse brain (top) or human 293Flp-In cells expressing RBM5-Flag (bottom) were subjected to sedimentation through glycerol gradients. The proteins detected in gradient fractions by immunoblotting are indicated on the right. Peak gradient regions collected for immunoprecipitation are indicated by blue and green-colored brackets. Positions of 5S, 18S and 28S sedimentation markers are also shown. (B) 293 Flp-In cells induced to express SF3A3-Flag protein under a range of Doxycycline concentrations. Proteins detected by immunoblotting in whole cell RIPA lysates are indicated on the right. (C) Glycerol gradient sedimentation of HMW extract proteins from 293Flp-In cells expressing SF3A3-Flag performed as in A bottom panel.

**Figure S2.**
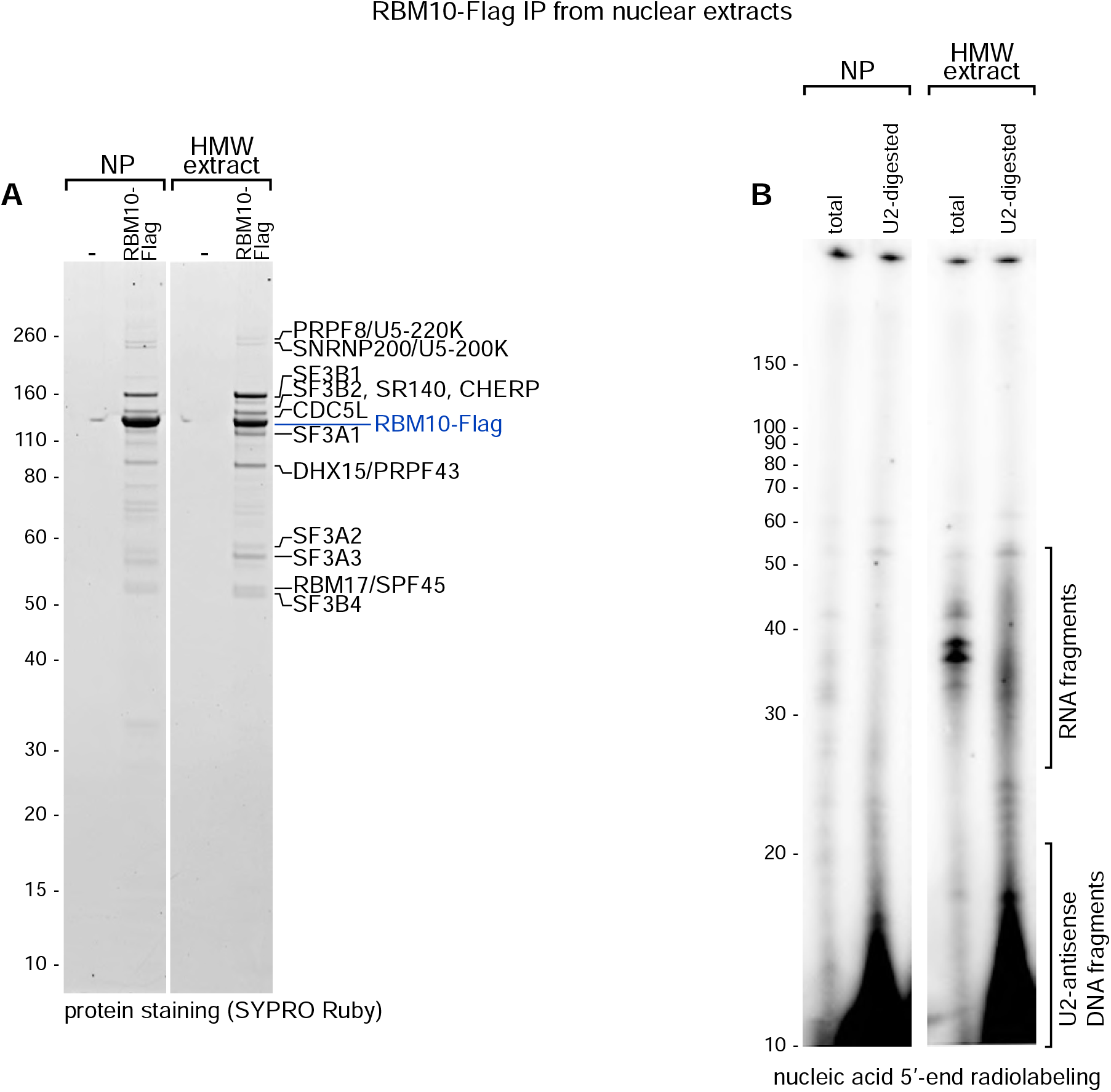
RBM10-Flag from HMW extract coprecipitates proteins and RNA fragments similar to those from the SF3A3 and RBM5 RNPs. (Related to Figure 1) (A) Protein profiles of anti-Flag-immunopurified material from nucleoplasm (lanes NP) and HMW nuclear extract from 293Flp-In cells not expressing Flag-tagged protein (lanes -) or expressing RBM10-Flag. (B) Analysis of coprecipitated RNA, performed as in Figure 1C.

**Figure S3.**
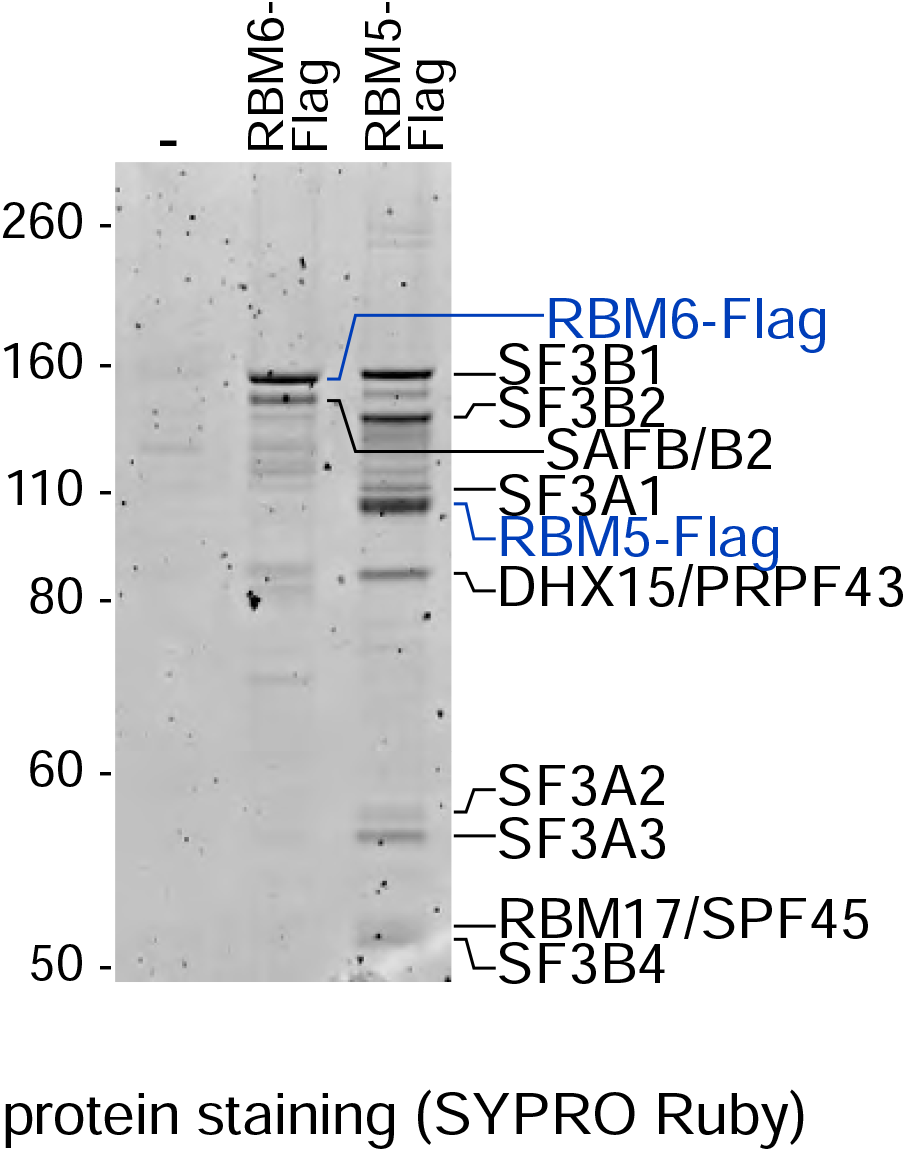
RBM6 does not coprecipitate with U2 snRNP components (Related to Figure 1) Comparison of protein profiles coprecipitated with RBM6-Flag or RBM5-Flag from 293Flp-In HMW nuclear extracts. Immunoprecipitation from cells not expressing a Flag-tagged protein serve as a negative control (lane -) The identities of the major protein bands are indicated on the right.

**Figure S4.**
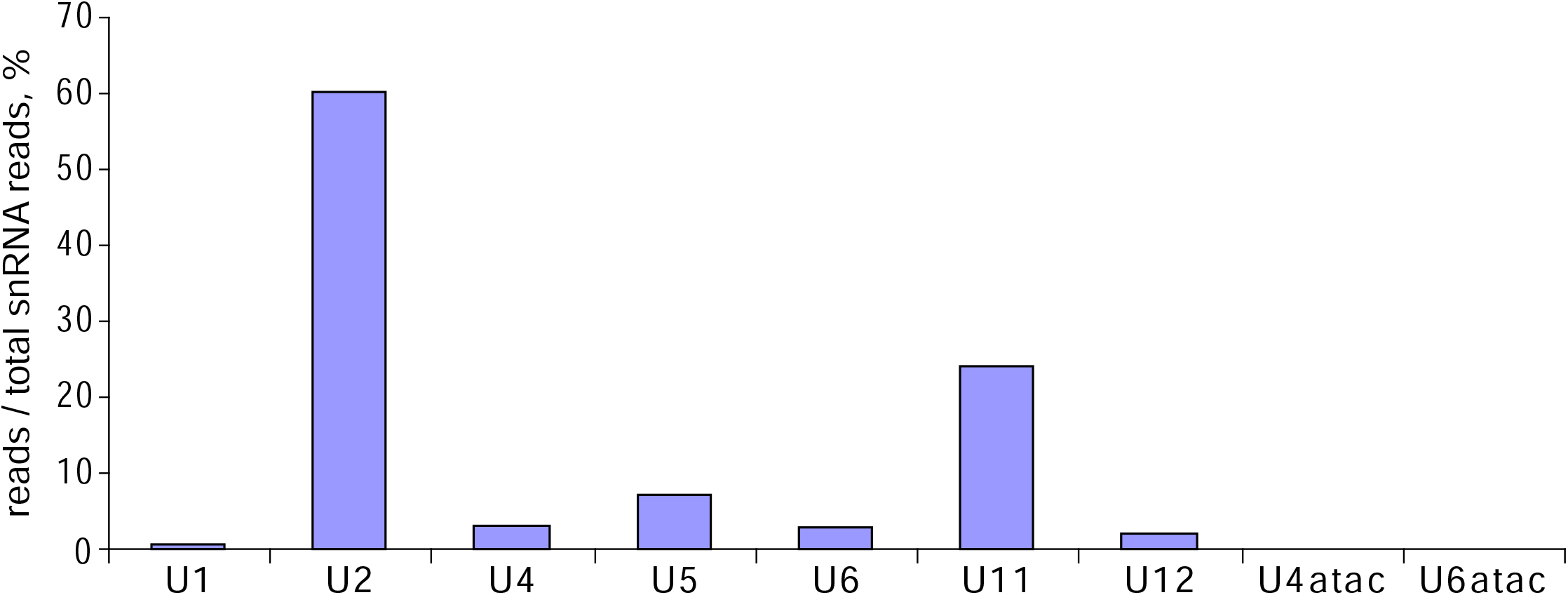
The RBM5 RNP complex contains U2 as well as U11/U12 snRNP components (Related to Figure 1 and Figure 2) Sequencing reads from the RBM5-Flag containing RNP (see Figure 1C lane total) were aligned to a library of human snRNA sequences. Unambiguous individual snRNA matches were counted and graphed as percentages of the total number of identified snRNA reads.

**Figure S5.**
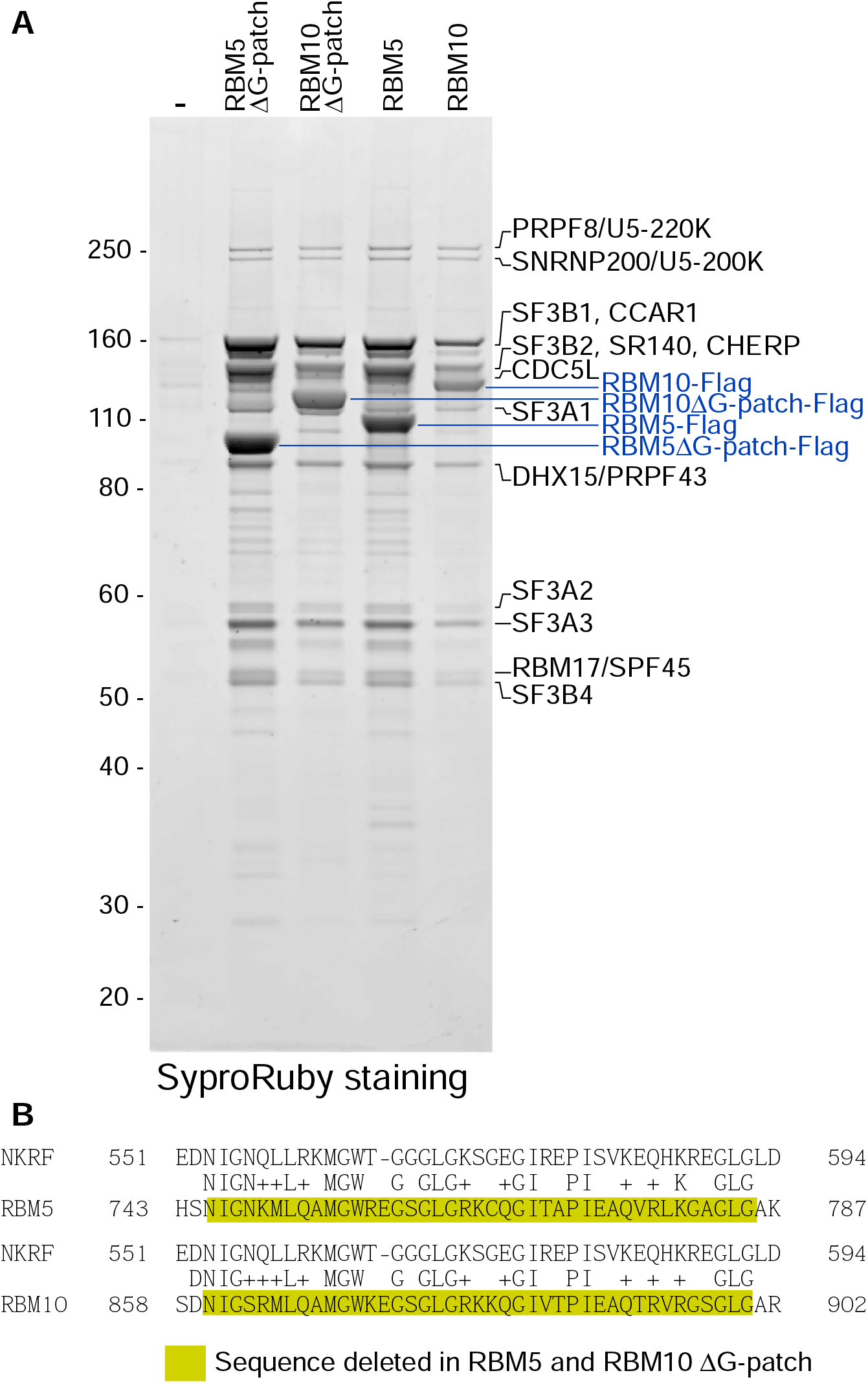
The G-patch motif of RBM5 and RBM10 is not required for interaction with U2 in the spliceosome (related to Figures 1 and 2). (A) Proteins coprecipitating with full-length or G-patch deletion mutants of RBM5 and RBM10 from HMW nuclear extracts. Extract from cells not expressing Flag-tagged protein (lane -) was also included. The immunoprecipitation is performed as in Figure S2A. The Flag-tagged proteins are indicated in blue and the major coprecipitated proteins in black. (B) Alignment of RBM5 and RBM10 G-patch motif sequences with NKRF, which interacts with DHX15 in pre-ribosomal complexes ^55^. The sequences deleted in A are highlighted in yellow.

**Figure S6.**
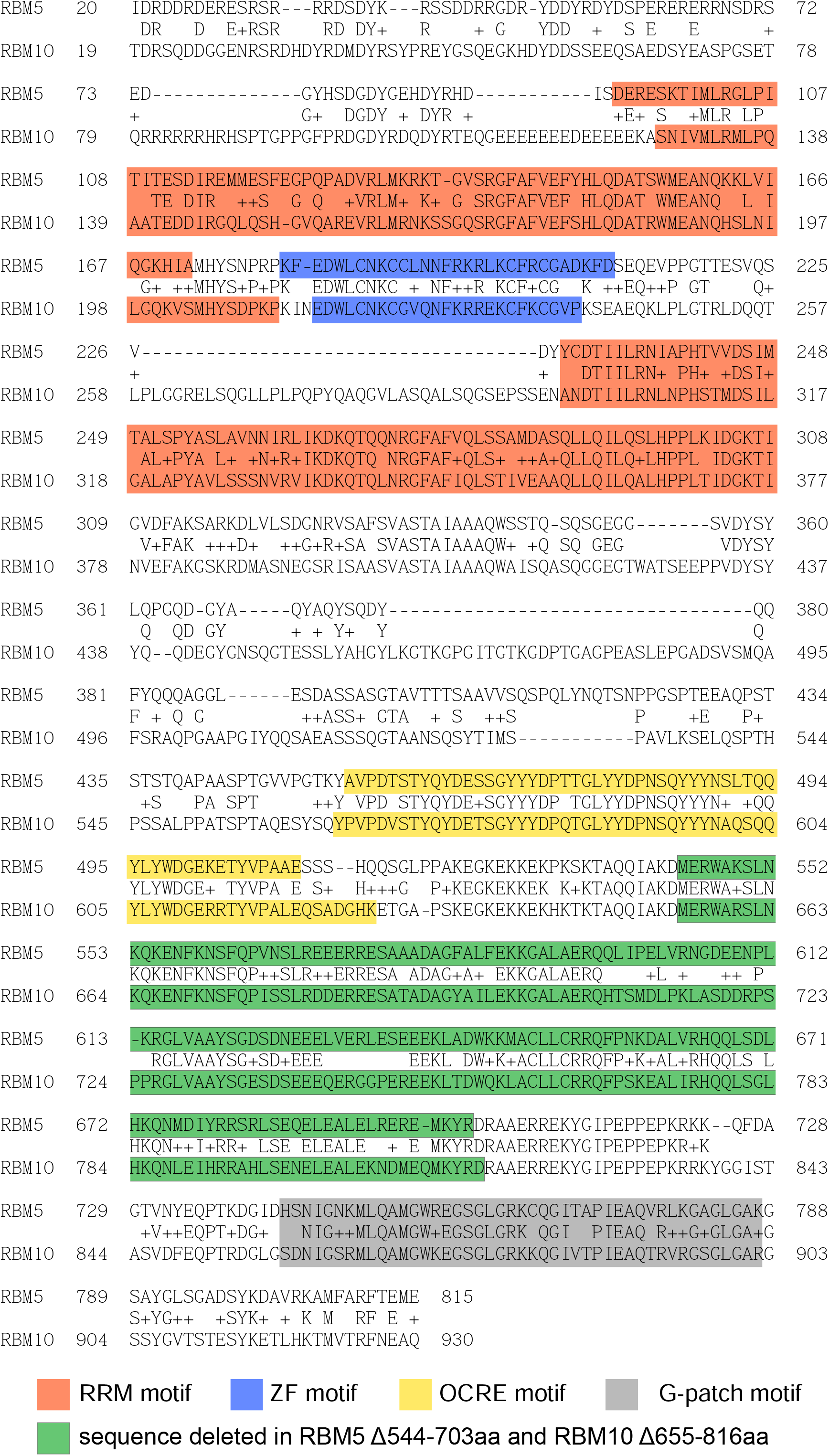
RBM5 and RBM10 share extensive protein sequence similarity and domain organization (Related to Figure 2). RBM5 and RBM10 proteins were aligned by BLAST ^62^, with known sequence motifs indicated ^68^. Deleted regions between the OCRE and the G-patch motifs of these proteins are shown in green.

**Figure S7.**
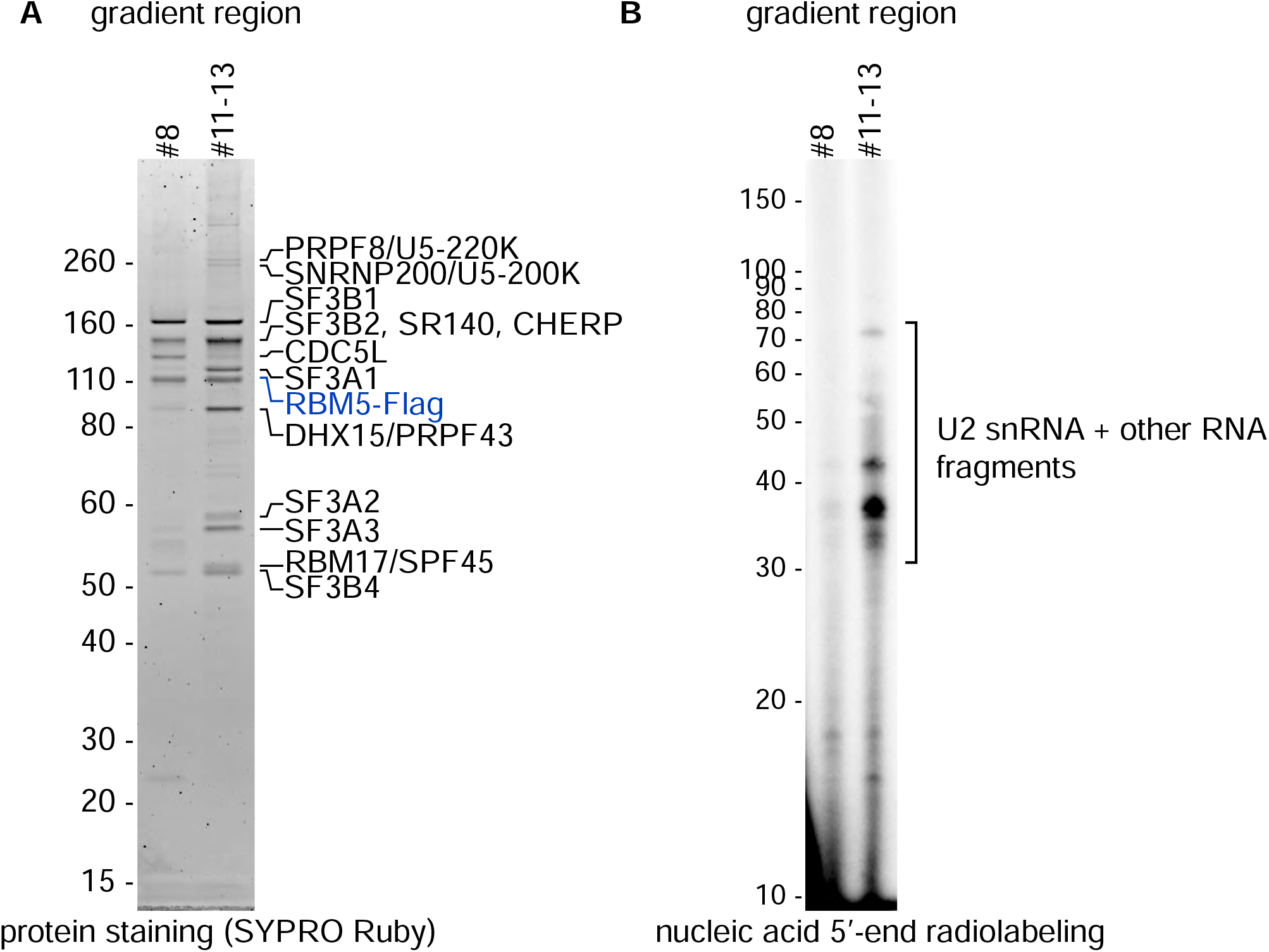
RBM5 interacts with a complex lacking SF3A. (Related to Figure 1 and Figure 2). RBM5-Flag RNP complexes from HMW extract were isolated from two separate regions of glycerol gradients as indicated in Figure S1A. Protein (A) and RNA analyses (B) of these complexes were performed as in Figure 1A and 1C, respectively.

**Figure S8.**
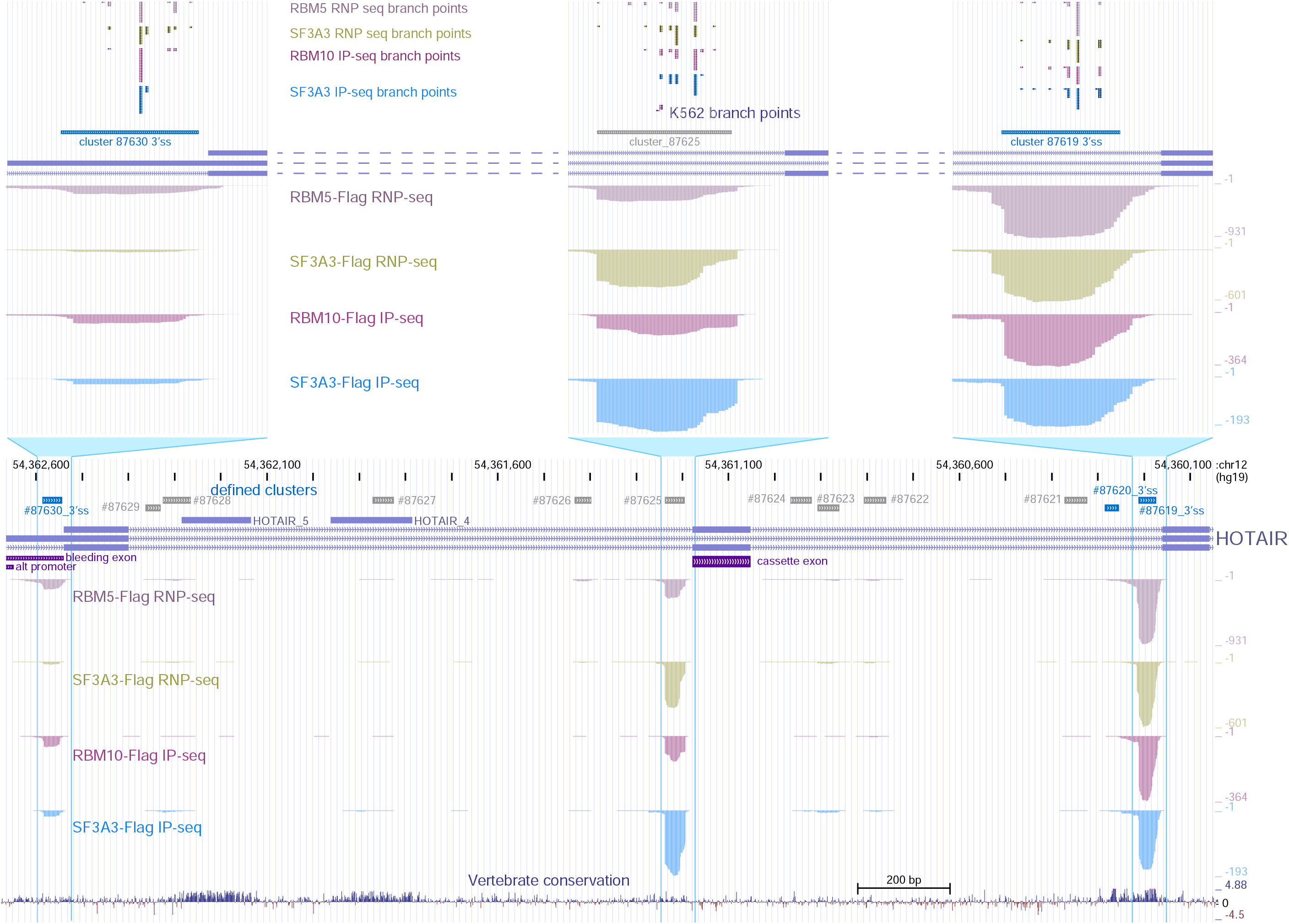
RBM5 and RBM10 RNPs are bound to branch sites of alternatively spliced and constitutive exons. (Related to Figure 3.) UCSC Genome browser view of protected RNA sequencing library reads mapped to the first three exons of the HOTAIR gene. Total read coverage is shown below the gene diagram (bottom), with individual RBP-seq or IP-seq tracks and clusters indicated as in Figure 3A. The tree prominent branch site clusters are shown in greater detail above. Previously identified and predicted precatalytic branch points are shown as in Figure 3B. See Figure S10 for details.

**Figure S9.**
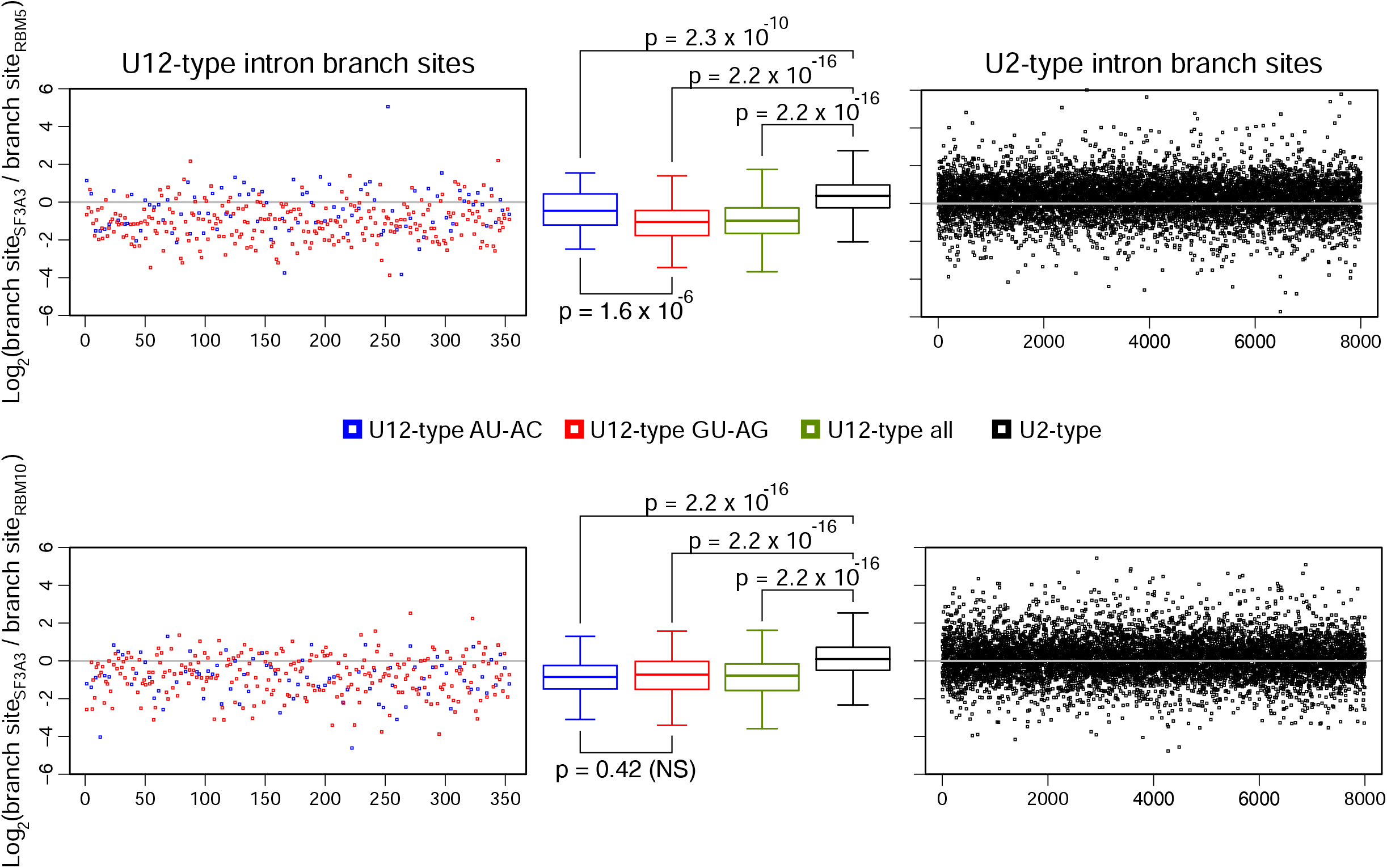
SF3A3 recovers reduced amounts U12-type branch sites compared to RBM5. (Related to Figure 3.) Comparison of RPKM ratios of SF3A3 over RBM5 RNP-seq reads (top) or over RBM10 IP-seq reads (bottom) from branch site clusters at annotated U12-type introns (left)^67^. The AU-AC and GU-AG intron subtypes are indicated in blue and red, respectively. A set of 8,000 randomly selected U2-type introns is similarly analyzed on the right in black. Between the U12 and U2 scatter plots, box plots of the median SF3A3 to RBM5 or RBM10 RPKM ratios for each intron group are shown on the same log_2_ scale with interquartile regions indicated. The median difference of each U12-type group of introns is compared to U2-type by Mann-Whitney U test with p-values indicated. NS indicates not significant.

**Figure S10.**
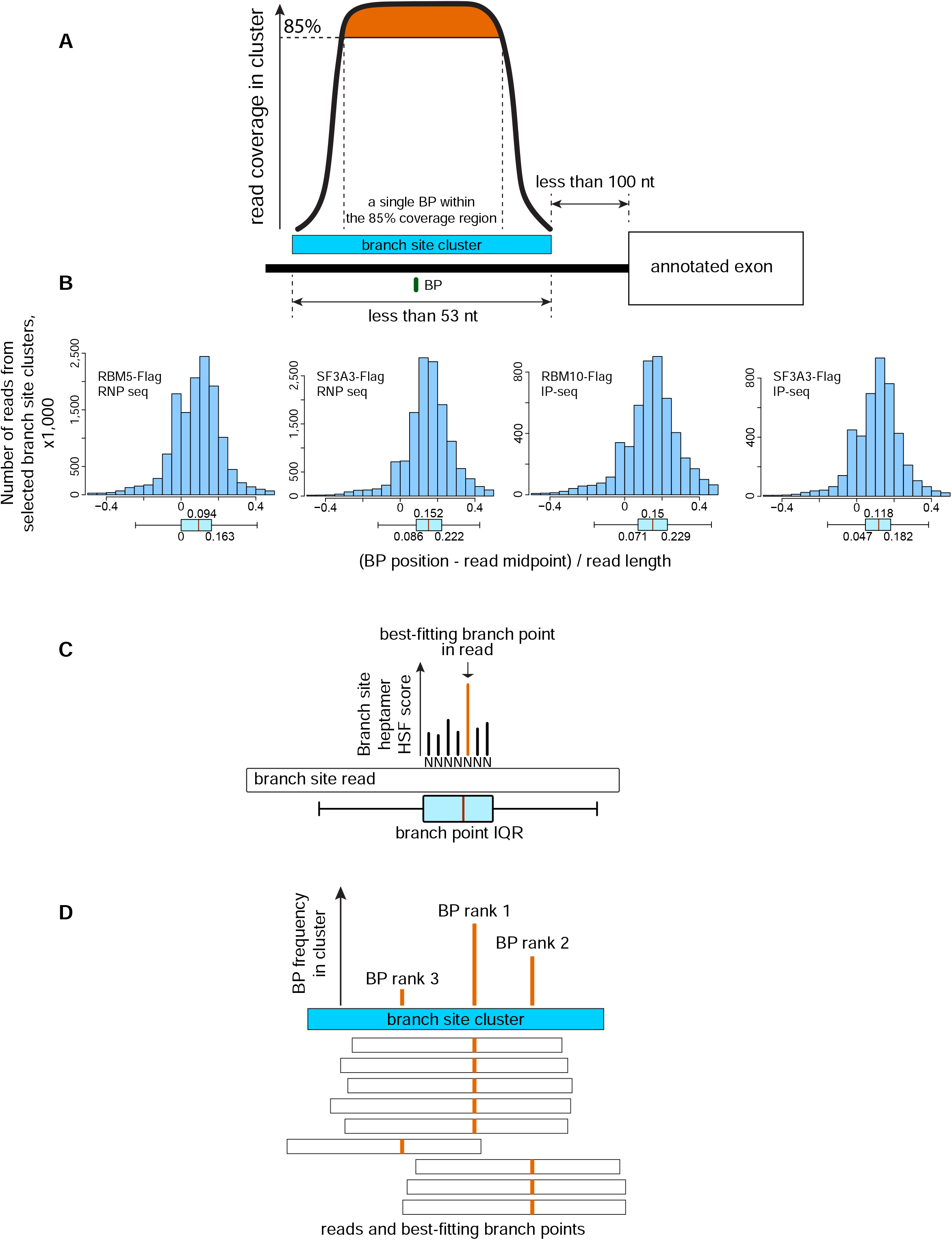
Prediction of branch point nucleotides within branch site clusters (Related to Figure 3). (A) Selection of a subset of approximately 29,000 clusters upstream of annotated exons and overlapping with a single identified branch point in previous studies ^37,38^, located within the region of highest read coverage for the cluster. (B) Distribution of branch point positions in reads from clusters selected in A. Box plots of the median branch point position with the interquartile ranges relative to the middle of the reads (position 0) are shown below for each RNP-seq and IP-seq dataset. (C) Identification of best-fitting the branch point in each read from all clusters as the nucleotide within the IQR interval with the highest branch site heptamer HSF score ^65,66^. (D) Ranking of the identified best-fitting branch points within a cluster.

**Figure S11.**
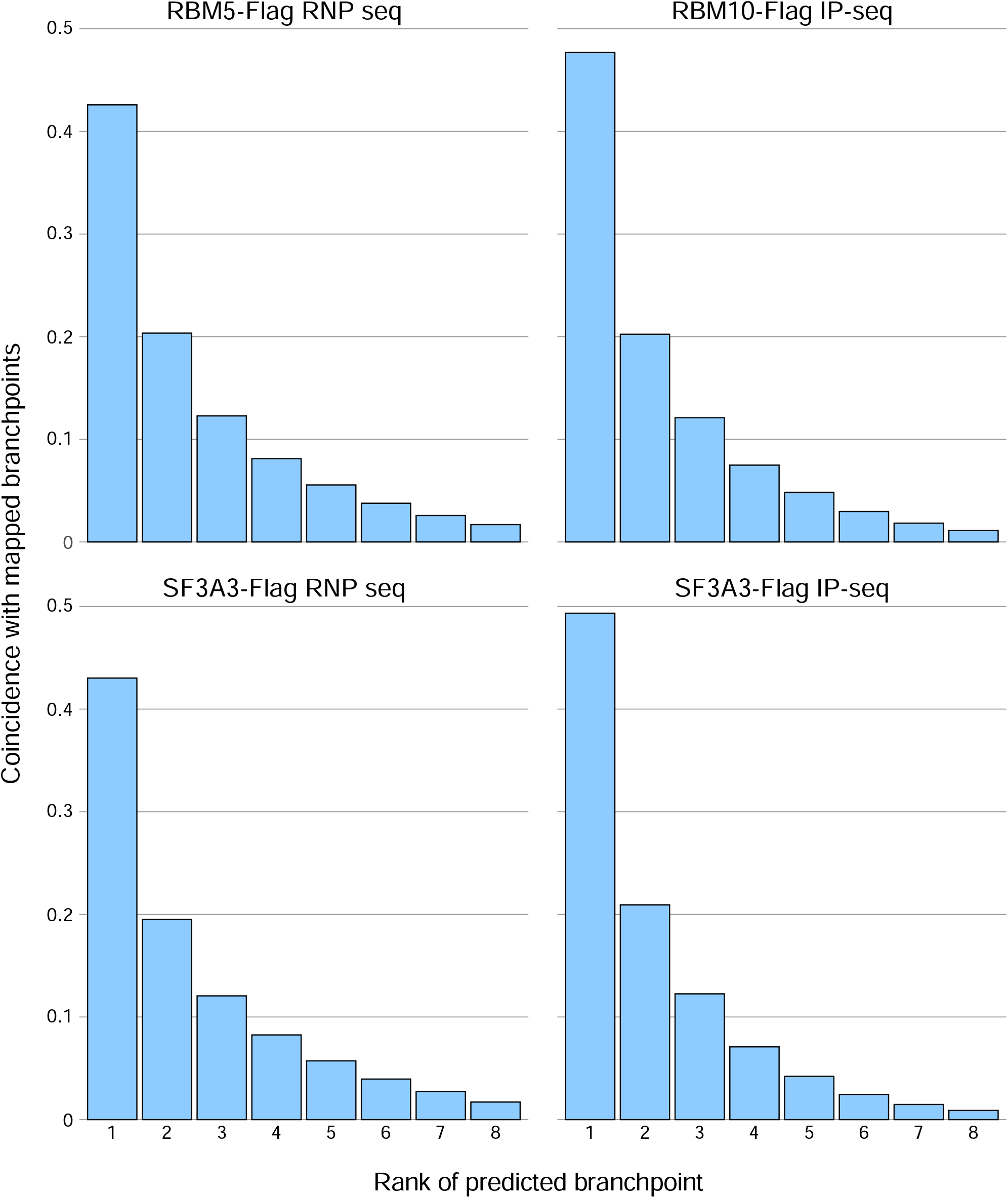
Coincidence of predicted and known branch points (Related to Figure 3). Fraction of predicted precatalytic branch points from RNP-seq and IP-seq clusters at intronic 3’ ends that align with published branch points (Mercer TR et al 2015, Zhang P et al 2022). The predicted branch points are grouped by their rank in each cluster.

**Figure S12.**
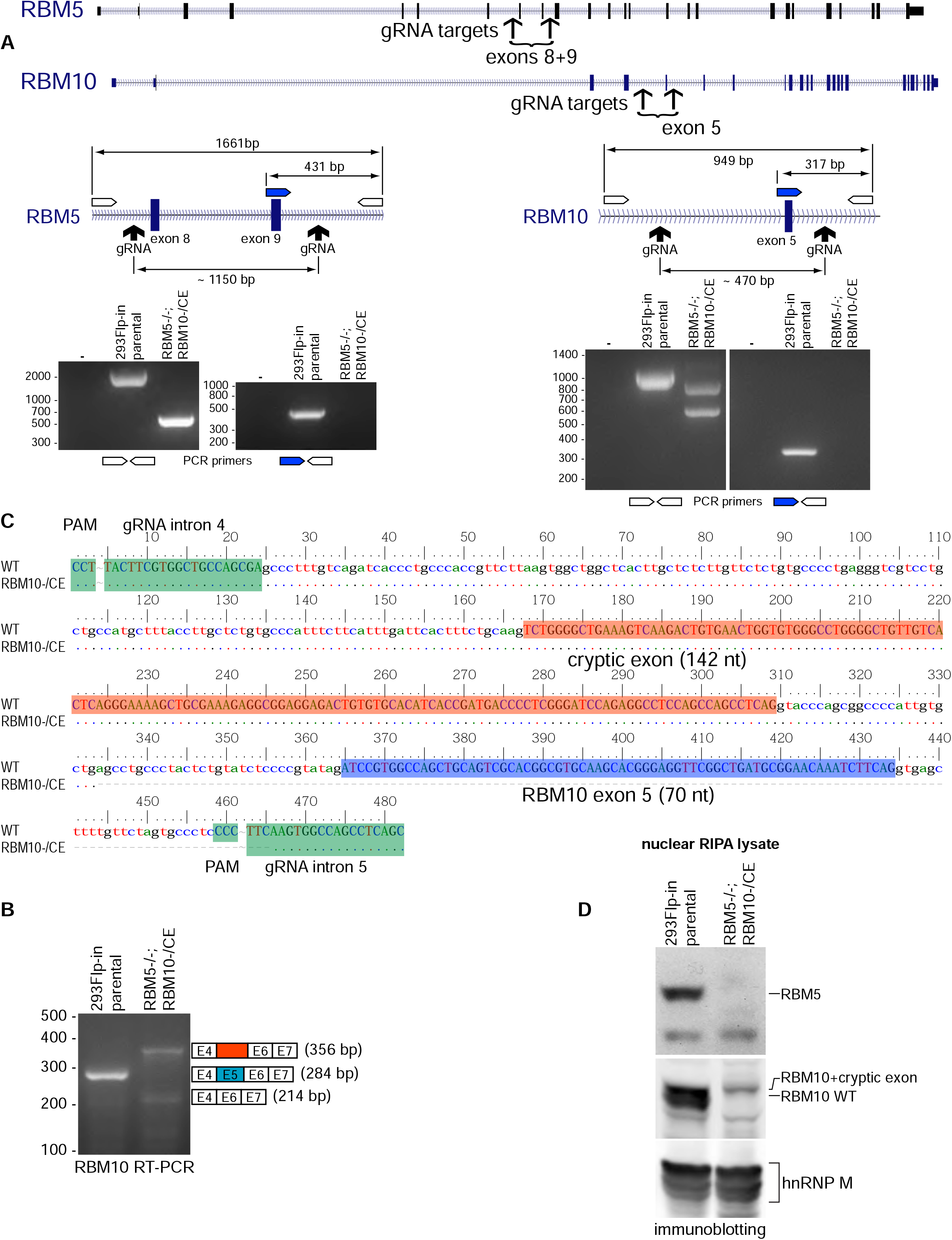
Genomic and gene expression characterization of 293Flp-In RBM5^-/-^; RBM10^-/CE^ cells (Related to Figure 4). (A) RBM5 and RBM10 gene structure diagrams. The positions of sgRNA targets used for gene editing, the genotyping primers, and the relevant distances are indicated (top). PCR of genomic DNA from parental 293Flp-In and RBM5^-/-^; RBM10^-/CE^ cells, with the indicated flanking or internal + flanking primer pairs. Lanes (-) indicate negative controls without DNA. The PCR products were resolved by agarose electrophoretic gel and stained with Ethidium bromide (bottom). (B) Analysis of RBM10 mRNAs produced in the edited clone. RT-PCR products from 293Flp-In and RBM5^-/-^; RBM10^-/CE^ cells using primers flanking RBM10 exons were resolved on agarose gel and stained with Ethidium bromide. The mRNA isoforms corresponding to detected bands are indicated on the right. (C) Characterization of the partially deleted RBM10 allele. Top band products from RBM5^-/-^; RBM10^-/CE^ genomic PCR and RT-PCR reactions were excised from gel, sequenced, and aligned to the genome. The positions of the RBM10 sgRNAs are indicated in green. Dashes below the sequence denote the region deleted in the sequenced RBM5^-/-^; RBM10^-/CE^ allele. RBM10 exon 5 is highlighted in blue and a cryptic exon partially included from this allele is shown in red. (D) RBM5 and RBM10 protein expression in parental and the RBM5^-/-^; RBM10^-/CE^ cells. Nuclear RBM5 and RBM10 were detected by immunoblotting. Wild type protein and RBM10 from the mutant allele containing the cryptic exon are indicated. hnRNP M immunoblotting serves as a loading control.

**Figure S13.**
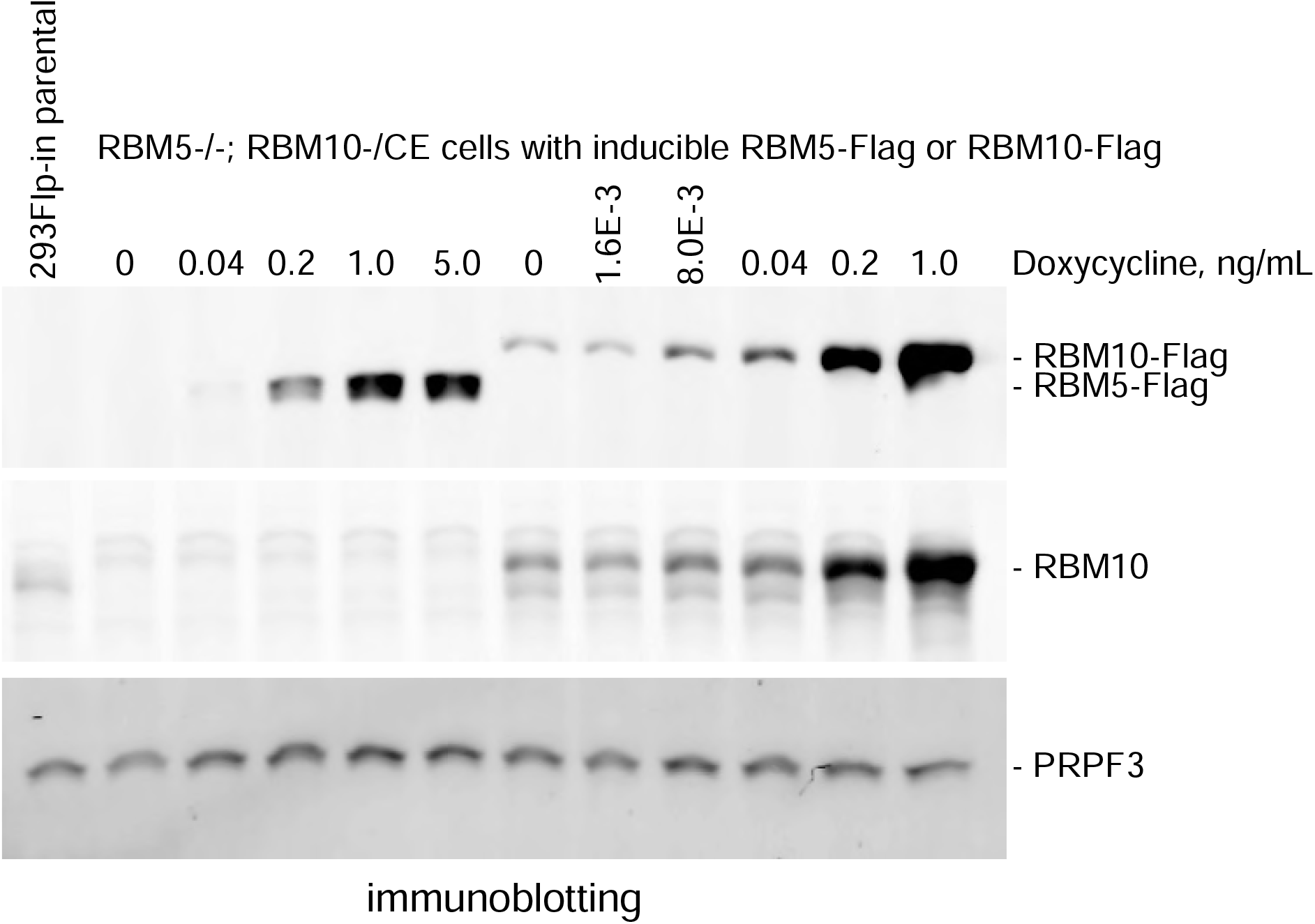
Comparison of RBM5-Flag and RBM10-Flag induction in RBM5^-/-^; RBM10^-/CE^ cells (Related to Figure 3). RBM5-Flag or RBM10-Flag transgenes were integrated into RBM5^-/-^; RBM10^-/CE^ cells at the Flip-in locus. These cells were induced to express the Flag-tagged proteins at the indicated Doxycycline concentrations. Protein levels were compared to parental 293Flp-In cells by immunoblotting from whole cell RIPA lysates. The detected proteins are indicated on the right.

**Figure S14.**
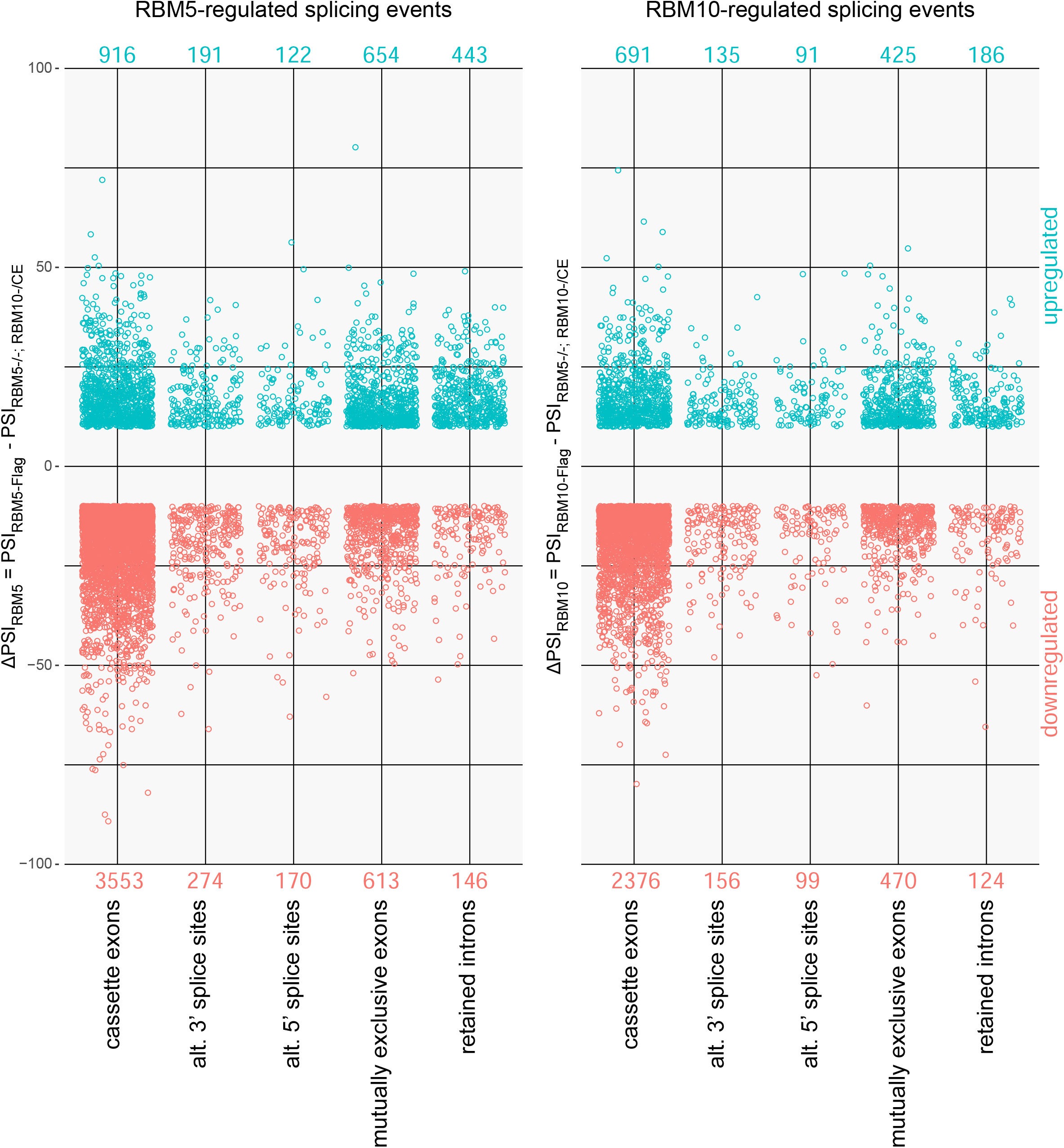
Significant RBM5-dependent and RBM10-dependent splicing events. (Related to Figure 4). RBM5 and RBM10 dependent splicing events were identified by rMATS as described in Figure 4AB. The number of upregulated and downregulated events from each type is shown on top or bottom, respectively. See also Tables S4 and S5.

**Figure S15.**
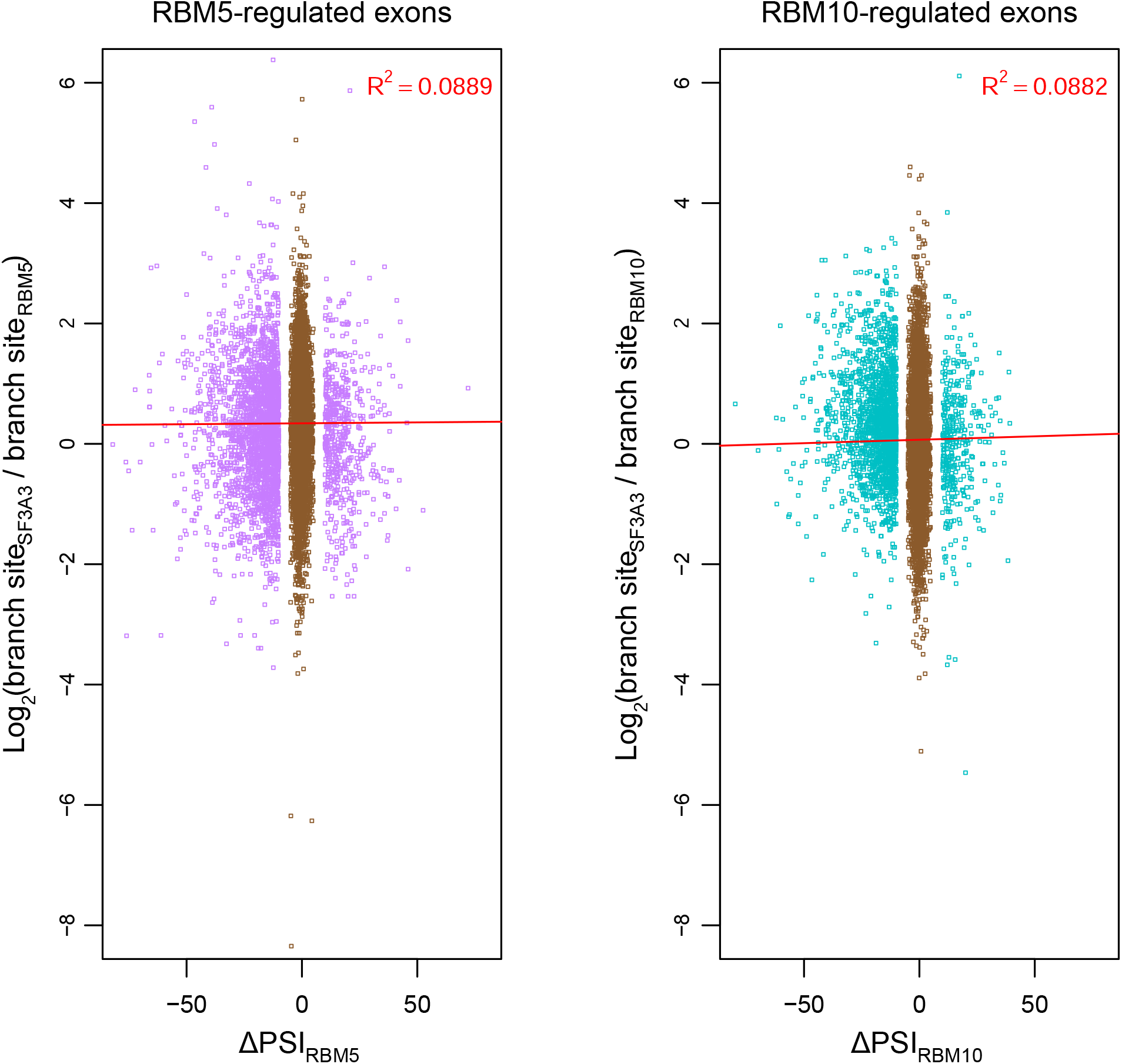
Branch site recovery by RBM5 RNP-seq and RBM10 IP-seq upstream of cassette exons does not correlate with exon regulation. (Related to Figure 4) Left: The ratio of RPKM values for SF3A3 and RBM5 RNP-seq in branch site clusters upstream of RBM5-regulated exons (purple) and control cassette exons (brown). X-axis plots the RBM5 ΔPSI; Y axis plots the log_2_ ratios of SF3A3 and RBM5-Flag branchpoint clusters of each exon. Right: Corresponding analysis of RBM10-regulated exons (green) and control cassette exons (brown) is shown on the right.

**Figure S16.**
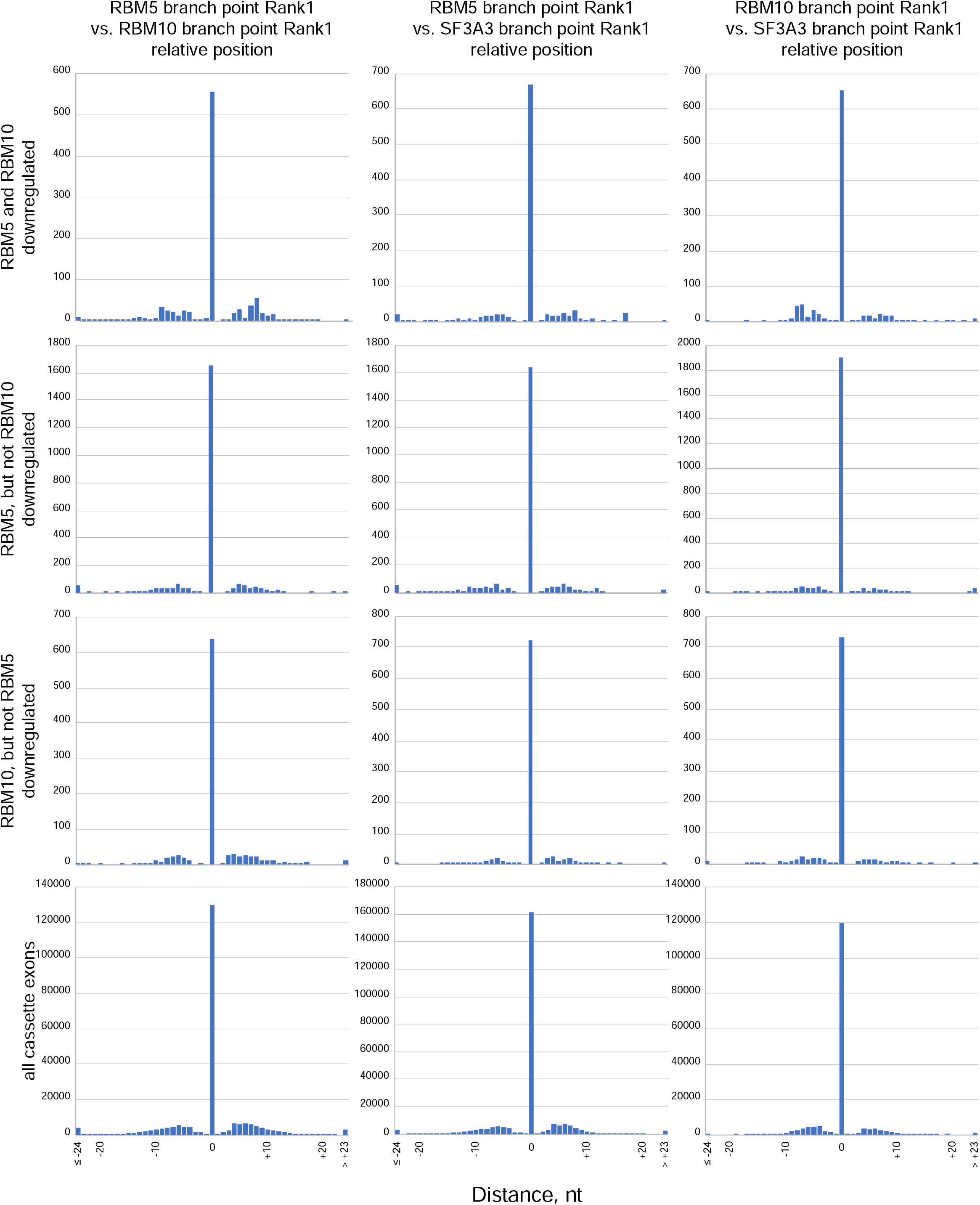
Predicted RBM5, RBM10, and SF3A3 precatalytic branch points overlap at similar rates upstream of regulated and unregulated exons. (Related to Figure 3 and Figure 4) Distances between the rank 1 branchpoints predicted in same clusters from the two RNP-seq or IP-seq datasets as indicated at the top. The nucleotide position of the branch point from the first dataset is set to 0. Negative or positive values on the X axis indicate the upstream or downstream location of the branch point from the second dataset. Y axis values are the number of branch site clusters exhibiting a deviation in branchpoint position between the compared groups. The subsets of cassette exons compared are indicated on the left.

**Figure S17.**
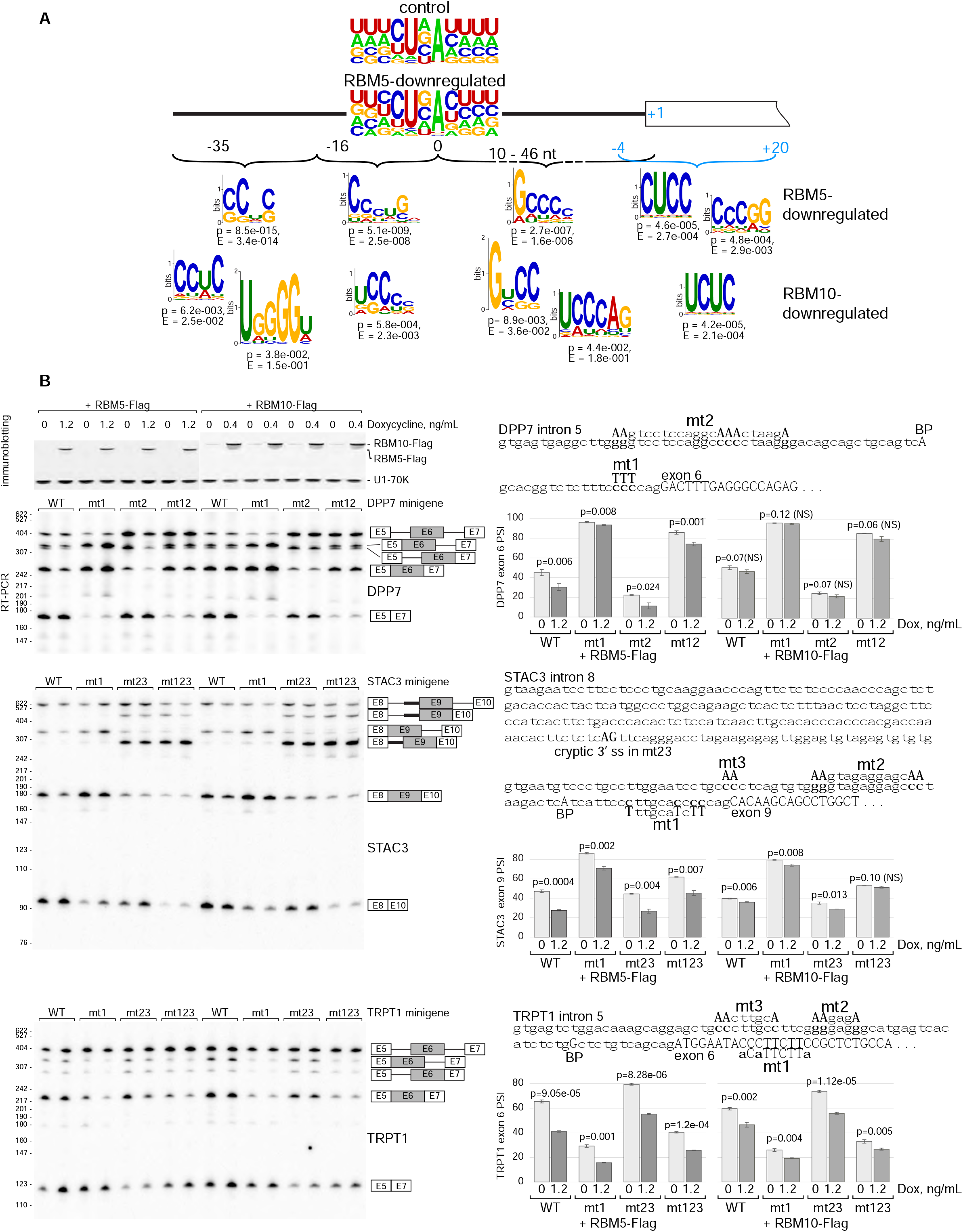
Sequence motifs enriched near the branch site of regulated exons. (Related to Figure 4.) (A) Diagram of a regulated exon with upstream intron indicating the major predicted RBM5 or RBM10 branch site as position 0 and the upstream and downstream intronic segments selected for motif enrichment analysis. A region including the last four intronic and the first 20 exonic nucleotides was also selected. These sequences were analyzed for motifs enriched relative to the corresponding regions in the control cassette exon set. The top motifs identified by STREME are shown below with enrichment scores and p values ^50^. WebLogo-generated branch site consensuses at RBM5-downregulated and control cassette exons are also shown ^69^. (B) Three-exon minigenes containing RBM5 and RBM10-downregulated cassette exons with full-length introns and flanking exons were transiently transfected in RBM5^-/-^; RBM10^-/CE^ cells containing RBM5-Flag or RBM10-Flag genes, and grown in presence or absence of Doxycycline. Flag-tagged protein expression is detected by immunoblotting (top). Splicing in wildtype (WT) and mutant minigenes is assayed 36 hours post-induction by RT-PCR using radiolabeled flanking primers followed by denaturing PAGE and phosphorimaging. One of three biological replicas is shown. Quantification of exon inclusion with error bars indicating standard deviations is also shown at the right. Significance was measured by T-test, with p-values shown above bar graphs. NS indicates not significant. Partial sequence of each minigene including the upstream intron in lowercase and cassette exon in uppercase is also shown on the right. Mutated nucleotides are shown in bold. BP indicates the major branch point nucleotide. Also shown in capitals is a cryptic 3’ splice site, active in STAC3 mt23 minigene.

**Fig. S18.**
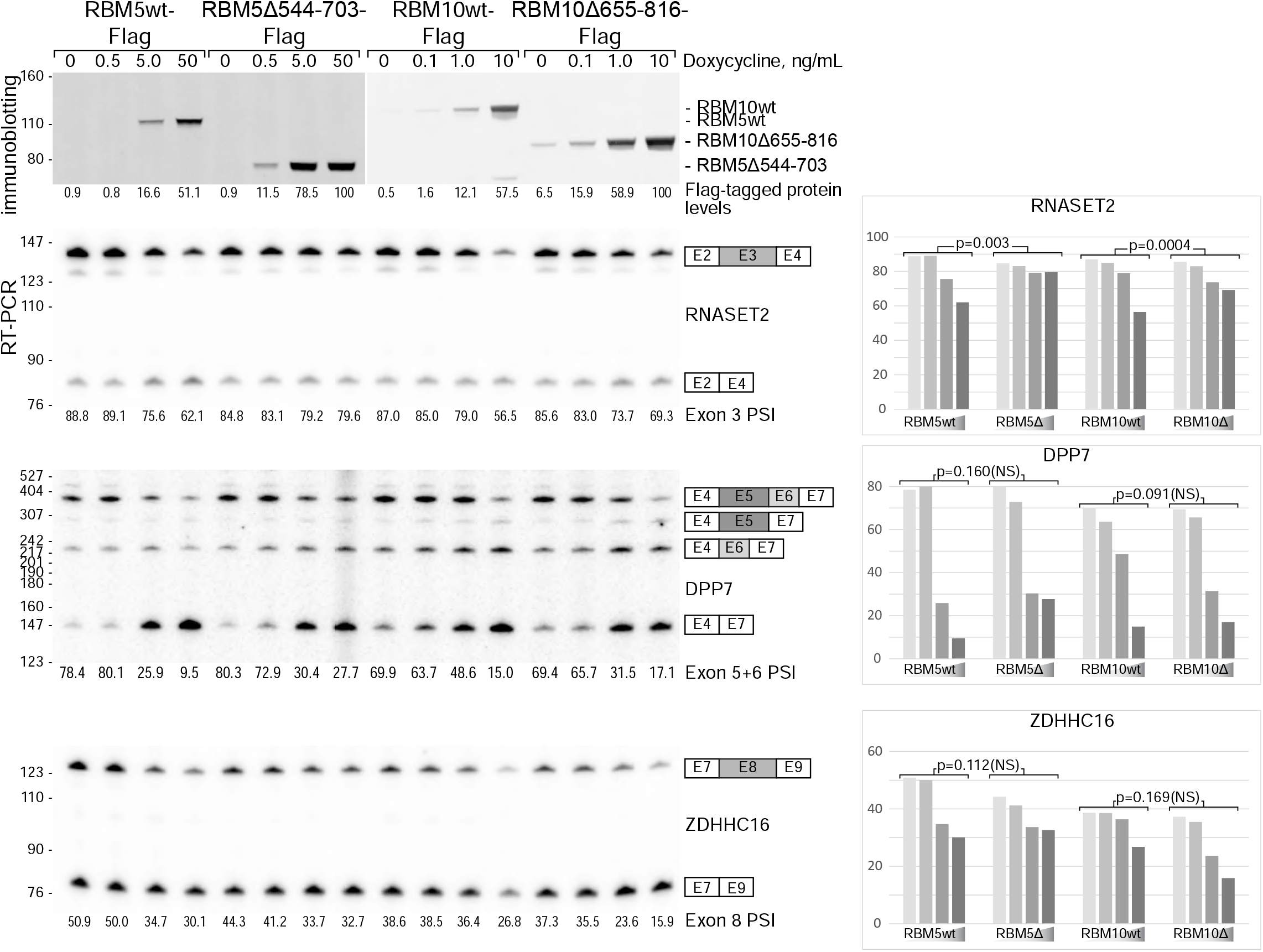
Splicing activity of RBM5 Δ544-703 and RBM10 Δ655-816 assayed on additional known downregulated exons. (Related to Figure 4) The immunoblot at the top is duplicated from Figure 4. The analysis these splicing events was the same as in Figure 4C.

## SUPPLEMENTARY TABLE LEGENDS

Table S1.

Mass-spectrometric analyses of samples immunoprecipitated from HMW extract with the indicated Flag-tagged protein. Proteins identified by LC-MS/MS and corresponding spectral counts are listed.

Table S2.

Mass-spectrometric analyses of samples immunoprecipitated with RBM5-Flag from the indicated glycerol gradient fractions of the HMW extract. Proteins identified by LC-MS/MS and corresponding spectral counts are listed.

Table S3. Summary of IP-seq and RNP-seq data. Numbers of genome-aligned reads and defined read clusters are listed for each of four conditions: RBM5 RNP-seq, RBM10 IP-seq, SF3A3 RNP-seq, and SF3A3 IP-seq. Numbers of these reads and clusters mapping to branch sites, 5’ splice sites, and to undefined genomic sites are also listed and their percentages.

Table S4. RBM5-dependent splicing changes

rMATS analyses splicing changes between RBM5^-/-^; RBM10^-/CE^ cells and these cells expressing ectopic RBM5-flag. Only significant events with |ΔPSI| > 0.1 and FDR < 0.1 are listed. For each splicing event, columns list gene name, exon coordinates, splice junction counts, average inclusion level for each condition, p value, FDR, and delta PSI. Individual tabs show data for alternative 3’ and 5’ splice sites, mutually exclusive exons, retained introns, and cassette exons.

Table S5. RBM10-dependent splicing changes

rMATS analyses splicing changes between RBM5^-/-^; RBM10^-/CE^ cells and these cells expressing ectopic RBM10-flag. Only significant events with |ΔPSI| > 0.1 and FDR < 0.1 are listed. For each splicing event, columns list gene name, exon coordinates, splice junction counts, average inclusion level for each condition, p value, FDR, and delta PSI. Individual tabs show data for alternative 3’ and 5’ splice sites, mutually exclusive exons, retained introns, and cassette exons.

