## Extended methods for "The apoptotic splicing regulators RBM5 and RBM10 are subunits of the U2 snRNP engaged with intron branch sites on chromatin"

### EXTENDED EXPERIMENTAL PROCEDURES

#### Antibodies

Primary antibodies used for immunoblot assays: FLAG (F3165-1MG, Sigma), DHX15 (Novus Biologicals, NBP2-13919), hnRNP M (Novus Biologicals, NB200-314), RBM5 (Thermo Fisher Scientific, PA5-35963), RBM10 (Thermo Fisher Scientific, PA5-83253), U2AF2 (Thermo Fisher Scientific, PA5-30442), PRPF3 (Thermo Fisher Scientific, PA5-96337), U2-A'/SNRPA1 (Abcam, ab128937), SNRPB (Smith antigen antibody Y12, Novus Biologicals, NB600-546), RBM17/SPF45 (Bethyl Laboratories, A302-498A). SF3A3: Rabbit monoclonal antibody against the C-terminal sequence KKTYEDLKRQGLL, U1-70K, SF3A1 and SF3B1 are described [1].

#### Minigene Sequences

TRPT1:
exon 5:
GCATGCGGTCCCATTGTGAAATAGCTGTGTTCATCGATGGACCCCTGGCTCTGGCAG
intron 5:
gtgagtctggacaaagcaggagctg**cc**cttgc**c**ttcg**gg**gag**g**gcatgagtcacatctc
**AA**cttgc**A** **AA**gag**A**
*branch point mt3 mt2
tgGctctgtcagcag
exon 6:
ATGGAATA**C**C**C**TTCTT**C**CGCTCTGCCAATGGGGTGATTCTGACTCCAGGGAATACTGAT
**a**C**a**TTCTT**a**
mt1
GGCTTCCTCCTTCCCAAGTACTTCAAGGAGGCCCTGCAGCTACGCCCTACCC
intron 6:
gtgagaaccaccaccccagcccctattccttgttcctgaaagctgtgcccttctgccct
cacctctctcctagcttccaaccctagactgacttaggtgtccctttctctaAccaact
cttgtctttatcag branch point*
exon 6:
GAAAGCCCCTTTCCTTGGCTGGTGATGAAGAGACAGAGTGTCAGAGTAGCCCCAAGCAC
AGCTCCAGAGAAAGGAGGAGGATCCAACAATAAAATATTAATTTATAAAAAAGAAATTT
TAAAAAGTAACAAGAAAGAACTCGTTTGAAACCATGTTTCATCATCCTGTA

STAC3:
exon 8:
GAAACCCTGAAGGGGATAAGAAGGCTGAGAAGAAGACACCTGATGACAAG
intron 8:

gtaagaatccttcctccctgcaaggaacccagttctctccccaacccagctctgacacc
actactcatggccctggcagaagctcactctttaactcctaggcttcccatcacttctg
acccacactctccatcaacttgcacacccacccacgaccaaaacacttctctcagttca
gggacctagaagagagttggagtgtagagtgtgtggtgaatgtccctgccttggaatcc
tgc**cc**ctcagtgtg**gg**gtagaggagc**cc**taagactcAtcattcc**c**ttgca**c**c**cc**cag
**AA** **AA**gtagaggagc**AA** * **T**ttgca**T**c**TT**
mt3 mt2 branch point mt1

exon 9:
CACAAGCAGCCTGGCTTCCAGCAGTCTCATTACTTTGTGGCTCTCTATCGGTTCAAAGC
CCTGGAGAAGGACGATCTGGATTTCCC

intron 9:
gtgagaggccttggggcaggggctggtactgcggagtgaagaggaaagagctgggttc
ccccaagggaaaggaaaagggggactcagggcttggagcctattgatgctggacccgg
ggcacagcaggggacctgctatgtgAgtgaggcccctcccttcccctcccag

exon 10: *branch point
GCCAGGAGAGAAGATCACAGTCATTGATGACTCCAATGAAGAATGGTGGCGG

DPP7:

exon 5:
TTATGGGGGATGCTCAGTGCCTACCTGAGGATGAAGTATCCCCACCTGGTGGCGGGG
CGCTGGCGGCCAGCGCGCCCGTTCTAGCTGTGGCAGGCCTCGGCGACTCCAACCAGTTC
TTCCGGGACGTCACGGCG

intron 5: mt2
**AA**gtcctccaggc**AAA**ctaag**A**
gtgagtgaggcttg**gg**gtcctccaggc**ccc**ctaag**g**gacagcagctgcagtcAgcacgg
tctctttc**ccc**cag branch point*
**TTT**
mt1

exon 6:
GACTTTGAGGGCCAGAGTCCCAAATGCACCCAGGGTGTGCGGGAAGCGTTCCGACAGAT
CAAGGACTTGTTCCTACAGGGAG

intron 6:
gtgaggtcctcccaccgcctcccactgcctgacccgccccagcgccagctatgcccaaa
cttcAagtctcTgccccttccacgcag
* *branch points

exon 7:
CCTACGACACGGTCCGCTGGGAGTTCGGCACCTGCCAGCCGCTGTCAGACGAGAAGGAC
CTGACCCAGCTCTTCATGTTCGCCCGGAATGCCTTCACCGTGCTGGCCATGATGGACTA
CCCCTACCCCACTGACTTCCTGGGTCCCCTCCCTGCCAACCCCGTCAAG

**Expression vectors**

RBM5-Flag, 2481bp-long sequence, inserted between the BamHI and XhoI sites of the vector pCDNA5/FRT/TO:

gccaccATGGGTTCAGACAAAAGAGTGAGTAGAACAGAGCGTAGTGGAAGATACGGTTCCATCATAGACAG
GGATGACCGTGATGAGCGTGAATCCCGAAGCAGGCGGAGGGACTCAGATTACAAAAGATCTAGTGATGATC

GGAGGGGTGATAGATATGATGACTACCGAGACTATGACAGTCCAGAGAGAGAGCGTGAAAGAAGGAACAGT
GACCGATCCGAAGATGGCTACCATTCAGATGGTGACTATGGTGAGCACGACTATAGGCATGACATCAGTGA
CGAGAGGGAGAGCAAGACCATCATGCTGCGCGGCCTTCCCATCACCATCACAGAGAGCGATATTCGAGAAA
TGATGGAGTCCTTCGAAGGCCCTCAGCCTGCGGATGTGAGGCTGATGAAGAGGAAAACAGGTGTAAGCCGT
GGTTTCGCCTTCGTGGAGTTTTATCACTTGCAAGATGCTACCAGCTGGATGGAAGCCAATCAGAAAAAGTT
GGTGATTCAAGGAAAGCACATTGCAATGCATTATAGCAATCCCAGACCTAAGTTTGAAGATTGGCTTTGTA
ACAAGTGCTGCCTTAACAATTTCAGGAAAAGACTAAAATGCTTCCGATGTGGAGCAGACAAGTTTGACTCT
GAACAGGAAGTGCCTCCTGGAACCACAGAGTCGGTTCAGTCTGTGGATTACTACTGTGATACGATCATTCT
TCGGAACATAGCTCCGCACACTGTGGTGGATTCCATCATGACAGCACTGTCTCCTTACGCGTCTTTAGCTG
TCAATAACATCCGCCTCATAAAAGACAAACAGACCCAGCAGAACAGAGGCTTCGCATTTGTGCAGCTGTCC
TCTGCAATGGATGCTTCTCAGCTGCTTCAGATATTACAGAGTCTCCATCCTCCTTTGAAAATTGATGGCAA
AACTATTGGGGTTGATTTTGCAAAAAGTGCCAGAAAAGACTTGGTCCTCTCAGATGGTAACCGCGTCAGCG
CTTTCTCTGTAGCTAGTACGGCTATTGCTGCTGCTCAGTGGTCATCCACCCAGTCTCAAAGTGGTGAAGGA
GGCAGTGTTGACTACAGTTATCTGCAACCAGGTCAAGATGGCTATGCCCAATATGCTCAGTATTCACAGGA
TTATCAGCAGTTTTATCAACAACAAGCTGGAGGATTGGAATCTGATGCATCATCTGCATCAGGCACAGCAG
TGACCACCACCTCAGCGGCTGTAGTGTCCCAGAGTCCTCAGCTGTATAATCAAACCTCCAATCCACCTGGC
TCTCCGACTGAGGAAGCACAGCCTAGCACTAGCACAAGTACACAGGCCCCAGCCGCTTCCCCTACTGGTGT
AGTTCCTGGTACCAAATATGCAGTACCTGACACGTCCACTTACCAGTATGATGAATCTTCAGGATATTACT
ATGATCCGACAACAGGGCTCTATTATGACCCCAACTCGCAATACTACTATAATTCCTTGACCCAGCAGTAC
CTTTACTGGGATGGGGAAAAAGAGACCTACGTGCCAGCTGCAGAGTCTAGCTCCCACCAGCAGTCGGGCCT
GCCTCCTGCAAAAGAGGGGAAAGAGAAGAAGGAGAAACCCAAGAGCAAAACAGCCCAGCAGATTGCCAAAG
ACATGGAACGCTGGGCTAAGAGTTTGAATAAGCAGAAAGAAAACTTTAAAAATAGCTTTCAGCCTGTCAAT
TCCTTGAGGGAAGAAGAAAGGAGAGAATCTGCTGCAGCAGACGCTGGCTTTGCTCTCTTTGAGAAGAAGGG
AGCCTTAGCTGAAAGGCAGCAGCTCATCCCAGAATTGGTGCGAAATGGAGATGAGGAGAATCCCCTCAAAA
GGGGTCTGGTTGCTGCTTACAGTGGTGACAGTGACAATGAGGAGGAGCTGGTGGAGAGACTTGAGAGTGAG
GAAGAGAAGCTAGCTGACTGGAAGAAGATGGCCTGTCTGCTCTGCCGGCGCCAGTTCCCGAACAAAGATGC
CCTAGTCAGGCACCAGCAACTCTCAGACCTTCACAAGCAAAACATGGACATCTATCGACGATCCAGGCTGA
GCGAGCAGGAGCTGGAAGCCTTGGAGCTAAGGGAGAGAGAGATGAAATACCGAGACCGAGCTGCAGAAAGA
CGGGAGAAGTACGGCATTCCAGAACCTCCAGAGCCCAAGCGCAAGAAGCAGTTTGATGCCGGCACTGTGAA
TTACGAGCAACCCACCAAAGATGGCATTGACCACAGTAACATTGGCAACAAGATGCTGCAGGCCATGGGCT
GGCGGGAAGGCTCTGGCTTGGGACGAAAGTGTCAAGGCATTACGGCTCCCATTGAGGCTCAAGTTCGGCTA
AAGGGAGCTGGCCTAGGAGCCAAAGGCAGCGCATATGGTTTGTCGGGCGCCGATTCCTACAAAGATGCTGT
CCGGAAAGCCATGTTTGCCCGGTTCACTGAGATGGAGATGGACTACAAAGACGATGACGACAAGtga

RBM5 Δ1-90aa: nucleotides 10-276 deleted.

RBM5 ΔRRM1+ZF: nucleotides 10-663 deleted.

RBM5 ΔRRM2: nucleotides 664-1068 deleted.

RBM5 ΔOCRE: nucleotides 1360-1554 deleted.

RBM5 Δ544-703aa: nucleotides 1636-2112 deleted.

RBM5dG-patch: deletion of nucleotides 2239-2361.

RBM5 ΔCT: nucleotides 2239-2451 deleted.

RBM10-Flag, 2823bp-long sequence, inserted between the HindIII and NotI sites of the vector pCDNA5/FRT/TO:

gccaccATGGAGTATGAAAGACGTGGTGGTCGTGGTGACAGGACTGGCCGCTATGGAGCCACTGACCGCTC
GCAGGATGATGGTGGGGAGAACCGCAGCCGAGACCACGACTACCGGGACATGGACTACCGTTCATATCCTC
GCGAGTATGGCAGCCAGGAGGGCAAGCATGACTATGACGACTCATCTGAGGAGCAGAGTGCGGAGGATTCC
TACGAGGCCTCCCCGGGCTCCGAGACTCAGCGTAGGCGGCGGCGGCGGCACAGGCACAGCCCCACCGGCCC
GCCAGGCTTCCCCCGAGACGGCGACTATCGGGACCAGGACTATCGGACCGAGCAAGGGGAGGAGGAGGAGG
AGGAGGAGGATGAGGAGGAGGAGGAGAAGGCCAGTAACATCGTCATGCTGAGGATGCTGCCACAGGCAGCC
ACTGAGGATGACATCCGTGGCCAGCTGCAGTCGCACGGCGTGCAAGCACGGGAGGTTCGGCTGATGCGGAA

CAAATCTTCAGGTCAGAGCCGGGGCTTCGCCTTCGTCGAGTTTAGTCACTTGCAGGACGCTACACGATGGA
TGGAAGCCAATCAGCACTCCCTCAACATCCTGGGCCAGAAGGTGTCGATGCACTACAGTGACCCCAAGCCC
AAGATCAATGAGGACTGGCTGTGCAATAAGTGTGGCGTCCAGAACTTCAAACGCCGAGAGAAGTGCTTCAA
ATGTGGCGTGCCCAAGTCAGAGGCAGAGCAGAAGCTGCCCCTCGGCACGAGGCTGGATCAGCAGACACTGC
CACTGGGTGGCCGGGAGCTGAGCCAGGGCCTGCTTCCCCTGCCGCAGCCCTACCAGGCCCAGGGAGTCCTG
GCCTCCCAAGCCCTGTCACAGGGCTCGGAGCCAAGCTCAGAGAACGCCAATGACACCATCATTTTGCGCAA
CCTGAACCCACACAGCACCATGGATTCCATCCTGGGGGCCCTGGCACCCTACGCGGTGCTGTCCTCCTCCA
ACGTGCGCGTCATAAAGGACAAGCAGACCCAACTGAACCGCGGCTTTGCCTTCATCCAGCTCTCCACCATC
GAGGCAGCCCAGCTGCTGCAGATCCTGCAGGCCCTGCACCCACCACTCACTATCGACGGCAAGACCATCAA
TGTTGAGTTTGCCAAGGGTTCTAAGAGGGACATGGCCTCCAATGAAGGCAGTCGCATCAGTGCTGCCTCTG
TGGCCAGCACTGCCATTGCTGCGGCCCAGTGGGCCATCTCACAGGCCTCCCAAGGTGGGGAGGGTACCTGG
GCCACCTCCGAGGAGCCGCCGGTCGACTACAGCTACTACCAACAGGATGAGGGCTATGGCAACAGCCAGGG
CACAGAGTCTTCCCTCTATGCCCATGGCTACCTCAAGGGCACCAAGGGCCCTGGCATCACTGGAACCAAAG
GGGATCCCACTGGAGCAGGTCCCGAGGCCTCCCTAGAGCCTGGGGCCGACTCTGTGTCGATGCAGGCTTTC
TCTCGCGCCCAGCCTGGTGCTGCTCCTGGCATCTACCAACAATCAGCCGAGGCGAGCAGTAGCCAGGGCAC
TGCTGCCAACAGCCAGTCGTATACCATCATGTCACCCGCTGTGCTCAAATCTGAGCTCCAGAGCCCTACCC
ATCCTAGTTCTGCTCTCCCACCGGCTACCAGCCCCACTGCCCAGGAATCCTACAGCCAGTACCCTGTTCCC
GACGTCTCTACCTACCAGTACGATGAGACCTCCGGCTACTACTATGACCCCCAGACCGGCCTCTACTATGA
CCCCAACTCCCAGTATTACTACAATGCTCAGAGCCAGCAGTACCTGTACTGGGATGGGGAGAGGCGGACCT
ATGTTCCCGCCCTGGAGCAGTCGGCCGACGGACATAAGGAGACAGGGGCACCCTCGAAGGAGGGCAAAGAG
AAGAAGGAGAAGCACAAGACCAAGACAGCTCAACAGATTGCCAAGGACATGGAACGCTGGGCCCGCAGTCT
CAACAAACAAAAAGAAAACTTCAAAAATAGCTTCCAGCCTATCAGCTCCCTGCGAGATGACGAGAGGCGGG
AGTCAGCCACTGCAGATGCTGGCTATGCCATCCTCGAGAAGAAGGGAGCACTAGCCGAGAGACAGCACACC
AGCATGGATCTCCCGAAATTGGCCAGTGACGACCGCCCAAGCCCTCCGCGAGGACTGGTGGCAGCCTACAG
CGGGGAGAGTGACAGTGAGGAGGAGCAGGAGCGTGGGGGCCCTGAGCGGGAGGAGAAGCTCACCGACTGGC
AGAAGCTGGCCTGTCTGCTCTGCCGACGCCAGTTCCCCAGCAAAGAGGCGCTCATCCGGCACCAGCAGCTC
TCAGGGCTCCACAAGCAAAACCTTGAGATTCACCGGCGcGCCCACTTGTCAGAAAACGAGCTAGAAGCACT
AGAGAAGAATGACATGGAGCAAATGAAGTACCGGGACCGTGCAGCTGAACGCAGAGAAAAGTATGGCATCC
CCGAGCCGCCAGAGCCCAAGAGGAGGAAGTACGGCGGCATATCCACAGCCTCTGTAGACTTCGAGCAGCCT
ACTCGGGACGGGCTGGGCAGTGACAACATTGGCAGTCGGATGCTGCAGGCCATGGGCTGGAAAGAGGGCAG
CGGCCTGGGCCGCAAGAAGCAGGGCATTGTAACGCCTATCGAGGCCCAAACACGGGTGCGGGGCTCCGGCC
TGGGTGCACGGGGCAGCTCCTACGGGGTCACCTCAACCGAGTCCTACAAGGAGACACTGCACAAGACAATG
GTGACCCGCTTCAACGAGGCCCAGATGGACTACAAAGACGATGACGACAAGtga

RBM10 Δ655-816aa: nucleotides 1966-2451 deleted.

RBM10 ΔG-patch: nucleotides 2584-2706 deleted.

SF3A3-Flag, 1535bp-long sequence, inserted between the BamHI and NotI sites of the vector pCDNA5/FRT/TO:

ccaccATGGAGACAATACTGGAGCAGCAGCGGCGCTATCATGAGGAGAAGGAACGGCTCATGGACGTCATG
GCTAAAGAGATGCTCACCAAGAAGTCCACGCTCCGGGACCAGATCAATTCTGATCACCGCACTCGGGCCAT
GCAAGATAGGTATATGGAGGTCAGTGGGAACCTGAGGGATTTGTATGATGATAAGGATGGATTACGAAAGG
AGGAGCTCAATGCCATTTCAGGACCCAATGAGTTTGCTGAATTCTATAATAGACTCAAGCAAATAAAGGAA
TTCCACCGGAAGCACCCAAATGAGATCTGTGTGCCAATGTCAGTGGAATTTGAGGAACTCCTGAAGGCTCG
AGAGAATCCAAGTGAAGAGGCACAAAACTTGGTGGAGTTCACAGATGAGGAGGGATATGGTCGTTATCTCG
ATCTCCATGACTGTTACCTCAAGTACATTAACCTGAAGGCATCTGAGAAGCTGGATTATATCACATACCTG
TCCATCTTTGACCAATTATTTGACATTCCTAAAGAAAGGAAGAATGCAGAGTATAAGAGATACCTAGAGAT
GCTGCTTGAGTACCTTCAGGATTACACAGATAGAGTGAAGCCTCTCCAAGATCAGAATGAACTTTTTGGGA
AGATTCAGGCTGAGTTTGAGAAGAAATGGGAGAATGGGACCTTTCCTGGATGGCCGAAAGAGACAAGCAGT
GCCCTGACCCATGCTGGAGCCCATCTTGACCTCTCTGCATTCTCCTCCTGGGAGGAGTTGGCTTCTCTGGG
TTTGGACAGATTGAAATCTGCTCTCTTAGCTTTAGGCTTGAAATGTGGCGGGACCCTAGAAGAGCGAGCCC
AGAGACTATTCAGTACCAAAGGAAAGTCCCTGGAGTCACTTGATACCTCTTTGTTTGCCAAAAATCCCAAG
TCAAAGGGCACCAAGCGAGACACTGAAAGGAACAAAGACATTGCTTTTCTAGAAGCCCAGATCTATGAATA
TGTAGAGATTCTCGGGGAACAGCGACATCTCACTCATGAAAATGTACAGCGCAAGCAAGCCAGGACAGGAG
AAGAGCGAGAAGAAGAGGAAGAAGAGCAGATCAGTGAGAGTGAGAGTGAAGATGAAGAGAACGAGATCATT

TACAACCCCAAAAACCTGCCACTTGGCTGGGATGGCAAACCTATTCCCTACTGGCTGTATAAGCTTCATGG
CCTAAATATCAACTACAACTGTGAGATTTGTGGAAACTACACCTACCGAGGGCCCAAAGCCTTCCAGCGAC
ACTTTGCTGAATGGCGTCATGCTCATGGCATGAGGTGTTTGGGCATCCCAAACACTGCTCACTTTGCTAAT
GTGACACAGATTGAAGATGCTGTCTCCTTGTGGGCCAAACTGAAATTGCAGAAGGCTTCAGAACGATGGCA
GCCTGACACTGAGGAAGAATATGAAGACTCAAGTGGGAATGTTGTGAATAAGAAGACATACGAGGATCTGA
AAAGACAAGGACTGCTCGACTACAAAGACGATGACGACAAGtga

**Primers**

Primers for RBM5 deletion genotyping:

RBM5 intron 7 forward: aactgtgtcccctgtccagt

RBM5 exon 8 forward: CCTCCTGGAACCACAGAGTC

RBM5 intron 9 reverse: atacacatgcattcccacca

Primers for RBM10 deletion genotyping:

hRBM10 intron 4 forward: aagcaatccccaggactacat

hRBM10 exon 5 forward: GCTGATGCGGAACAAATCTT

hRBM10 intron 5 reverse: tcagagtcccagagaggaggt

RT-PCR primers for RBM10 cryptic exon detection and sequencing:

hRBM10 exon 4 forward: GGCCAGTAACATCGTCATGC

hRBM10 exon 7 reverse: TATTGCACAGCCAGTCCTCA

RT-PCR primers for detection of RBM5 and RBM10-dependent cassette exons expressed from endogenous genes and three-exon minigenes:

TRPT1 exon 5 forward: CATGCGGTCCCATTGTGAAA

TRPT1 exon 7 reverse: TTCTCTGGAGCTGTGCTTGG

STAC3 exon 8 forward: CCTGAAGGGGATAAGAAGGCT

STAC3 exon 10 reverse: CGCCACCATTCTTCATTGGAGT

ZDHHC16 exon 7 forward: AGCTATGGAAGTTGGGACCT

ZDHHC16 exon 9 reverse: GAAAGGAGAAGGTGGGTGGT

RNASET2 exon 2 forward: ACTAATTATGGTTCAGCACTGGC

RNASET2 exon 4 reverse: AAATTGAAGGGCCACGATCT

NUMB exon 8 forward: TCTGCTCCGATGACCAAACC

NUMB exon 10 reverse: GTACGTCTATGACCGGCCTG

HOTAIR exon 2 forward: CATTCTGCCCTGATTTCCGG

HOTAIR exon 4 reverse: AATCCGTTCCATTCCACTGC

PCBP2 exon 11 forward: TGGCAATGCAACAGTCTCAT

PCBP2 exon 13 reverse: TCGTTTGGAATGGTGAGTTCA

DPP7 exon 4 forward: CCATCGCCTTCGGTGGAA

DPP7 exon 7 130-111: GTGGGGTAGGGGTAGTCCAT

(for endogenous gene splicing analysis)

DPP7 exon 5 forward: GCGACTCCAACCAGTTCTTC

DPP7 exon 7 reverse: CAGGAAGTCAGTGGGGTAGG

(for three-exon minigene splicing analysis)

**IPseq, RNPseq protocol**

This method used library construction similar to that described for iCLIP and shares some iCLIP reagents [2].

**Materials:**

**Tubes and tips:**

RNase-free 0.2 ml, 1.5 ml, 5 ml, 15 ml, and 50 ml tubes. Use barrier tips in all library construction steps, use separate pipette sets in pre- and post- PCR amplification steps.

**Oligonucleotides:**

**U2 antisense oligonucleotide**:

GGGTGCACCGTTCCTGGAGGTACTGCAATACCAGGTCGATGCGTGGAGTGGACGGAGCA
AGCTCCTATTCCATCTCCCTGCTCCAAAAATCCATTTAATATATTGTCCTCGGATAGAG
GACGTATCAGATATTAAACTGATAAGAACAGATACTACACTTGATCTTAGCCAAAAGGC
CGAGAAGCGAT

**L3 linker (RNA)**: /Phos/-UGAGAUCGGAAGAGCGGUUCAG-Biotin

In this method the 3' hydoxyl end may be blocked by modifications other than biotinylation.

**Primers for reverse transcription**:

**RTclipGTT**: /Phos/-nnnnAACnnnnAGATCGGAAGAGCGTCGTggatccTGAACCGC

**RTclipGCC**: /Phos/-nnnnGGCnnnn...

**RTclipATC**: /Phos/-nnnnGATnnnn...

*The barcodes are underlined. Order RT primers with different barcodes in such a way that no single nucleotide substitution should convert one barcode to another.*

**cut_oligo**: GTTCAGGATCCACGACGCTCTTCaaaa

**P5solexa**:

AATGATACGGCGACCACCGAGATCTACACTCTTTCCCTACACGACGCTCTTCCGATCT

**P3solexa**:

CAAGCAGAAGACGGCATACGAGATCGGTCTCGGCATTCCTGCTGAACCGCTCTTCCGATCT

**Procedure:**

**1. Preparation of samples:**

Homogenize cells, pellet nuclei through sucrose cushion, rinse and pellet nuclei as described [3,4]. Lyse pelleted nuclei for 5 minutes in 10 volumes of ice-cold lysis buffer (20 mM HEPES-KOH pH 7.5, 150 mM NaCl, 1.5 mM MgCl2, 0.5 mM CaCl2, 0.5 mM DTT, 1.25x Complete Protease Inhibitor Cocktail (Millipore-Sigma), and 0.6% Triton X-100). Pellet the nuclear HMW material at 20,000x g for 5 min at 4 °C. Remove the soluble nucleoplasm, and transfer into a fresh tube to process in parallel if needed. Add equal volume of lysis buffer to the HMW pellet. Adjust the volume of all samples to at least 400 µl. Add 0.04 U/µl TURBO DNase (Thermo Fisher Scientific) to HMW samples, and 5 U/µl of Benzonase Nuclease (Millipore-Sigma), 0.01 U/µl RNase A, and 0.4 U/µl RNase T1 (RNase Cocktail, Thermo Fisher Scientific) to all samples. Glycerol in samples, added from enzyme storage buffers, should not exceed 5% (v/v). Incubate at room temperature (22-24°C) on a rotator to digest RNA and DNA as described, then clear the extracted material by centrifugation for 10 min at 20,000x g, 4 °C [3,4]. Prepare 11.5 ml 10-30% glycerol gradients in 14x89mm ultra-clear ultracentrifugation tubes (Beckman Coulter), containing 20 mM HEPES-KOH pH 7.5, 150 mM NaCl, 1.5 mM MgCl2, 0.5 mM DTT, and 1x Complete Protease Inhibitor Cocktail. Overlay 250-400 µl of nucleoplasm or HMW extract on top, and centrifuge in SW41Ti rotor (Beckman Coulter) for 17 hours at 32,000 g, 4°C. Transfer desired gradient regions in appropriately-sized Slide-A-Lyzer MINI Dialysis Devices, 10K MWCO Membrane (Thermo Fisher Scientific). Dialyze twice for 30 min at 4°C with buffer containing 20 mM HEPES-KOH pH 7.5, 150 mM NaCl, 1.5 mM MgCl2, and 0.5 mM DTT. Incubate dialyzed samples (RNPseq) or HMW extracts (IPseq) with 5-7.5 µl packed-volume M2 FLAG agarose beads (Millipore-Sigma), rotating overnight at 4 ºC. Wash agarose beads four times with ice-cold wash buffer (20 mM HEPES-

KOH pH 7.5, 150 mM NaCl, 1.5 mM MgCl2, and 0.05% Triton-X100). Pellet the washed beads in swinging-bucket rotor centrifuge to minimize the loss of beads. After the last washing step and buffer removal, centrifuge the beads again in fixed-angle rotor for 30 sec at 20,000 xg, and remove the residual wash buffer with 10-μl tip. Elute for two hours at 4°C in 35-60 μl of buffer containing 20 mM HEPES-KOH pH 7.5, 150 mM NaCl, 1.5 mM MgCl2, and 150 ng/μl 3xFLAG peptide (Millipore-Sigma), shaking for 15 sec every 5 min in a Thermomixer (Eppendorf). Use 10-40% of the eluted material for protein analysis by SDS-PAGE and protein staining.

**2. Deproteinization:**

Incubate the eluted material in a 200-μl reaction containing 50 mM Tris-HCl pH 8.0, 5 mM EDTA, 0.5% SDS, and 5 U of Proteinase K (New England BioLabs) for 30 min at 55°C. Add an equal volume of Acid phenol:chloroform:isoamyl alcohol (125:24:1), pH 4.3–4.7 (Thermo Fisher Scientific) and vortex for at least 30 seconds. Centrifuge for 5 min at 20,000 xg, room temperature. Transfer the upper aqueous phase into a fresh tube and precipitate overnight in presence of 70% ethanol, 85 mM sodium acetate pH 5.4, and 15 μg of GlycoBlue (Thermo Fisher Scientific). Centrifuge for 20 min at 20,000 xg, 4°C, wash with 1 ml of 75% ethanol, and dry the pellet at room temperature. Dissolve the pelleted RNA in 6 μl of DEPC-treated MilliQ-water (Millipore-Sigma) and store at -20°C.

**3. U2 snRNA degradation:**

This step may be optionally skipped, or a portion of the sample saved for generation of sequencing libraries including recovered U2 snRNA. When skipped, adjust the volume to 20 μl with DEPC-treated water and proceed with step 4.

Mix 6 μl dissolved RNA with 2 μl of 100 ng/μl U2 antisense oligonucleotide containing 0.4 mM EDTA. Incubate for 5 min at 55°C in a thermal cycler. Adjust the temperature to 37°C, add the following premixed solutions: 1.5 μl of 5U/μl RNase H (New England BioLabs), 2.0 μl of the supplied 10x reaction buffer, 1.0 μl of 40 U/µl RNaseOUT (Thermo Fisher Scientific), 7.5 µl of DEPC-treated water. Incubate for 20 min at 37°C.

**4. Dephosphorylation and DNA degradation:**

Premix the following solution: 5 µl of 2 U/µl TURBO DNase, 20 µl of the supplied reaction buffer, 5 µl of 1U/µl FastAP alkaline phosphatase (Thermo Fisher Scientific), 2.5 µl of 40 U/µl RNaseOUT, and 147.5 µl DEPC-treated water. Mix this solution with the RNase H reaction, and incubate for additional 45 min at 37°C. Add 2 µl of 500 mM EDTA, and perform phenol extraction and precipitation as in step 2, omitting the addition of GlycoBLue. Dissolve the dried pellet in 5.5 µl of DEPC-treated water and store at -20°C.

**5. Denaturing PAGE analysis of the recovered RNA fragments:**

This is an optional step to assess the quantity and the size of the recovered RNA. Add 0.5 µl of dephosphorylated RNA per 7-µl reaction, also containing 3.33 U of T4 Polynucleotide Kinase (New England BioLabs), 1x supplied reaction buffer, 8 U of RNaseOUT, and approximately 100 µCi of γ-[$^{32}$P] ATP (PerkinElmer). Incubate for 15-20 min at 37°C, add 10 µl of denaturing buffer (formamide containing 10 mM EDTA pH 8.0, and Bromophenol Blue Xylene Cyanole Dyes at appropriate color density). Incubate at 85 °C for 5 min, and immediately place on ice for at least one minute. Resolve on 10% acrylamide (Acrylamide:Bisacrylamide, 19:1), 50% urea, 1x TBE gel after electrophoresis pre-run for at least 10 minutes. Resolve MspI-digested pBR322, or other suitable radiolabeled marker along samples. Dry the gel and image by phosphorimaging on Amersham Typhoon (GE Healthcare Bio-Sciences).

**6. Size-selection of recovered RNA:**

Mix 5 µl of RNA solution with 7.5 µl of denaturing buffer. Mix GeneScan 500 and 600 LIZ fluorescent markers (Thermo Fisher Scientific) with denaturing buffer not containing dies. Denature samples and markers, load markers corresponding to 0.5-1.0 µl of stock solution per lane on either side of the samples. Resolve as in

step 5 on a 5.5% acrylamide gel. Electrophoresis should be run for a shorter time so the in-gel distance between the lower and the upper-size limits is about 10 mm. Rinse the glass gel plates with distilled water and wipe until dry, but do not disassemble. Scan the gel on Amersham Typhoon in the Cy5 channel to detect the bands of the marker. Print real and mirror images in actual size, draw excision guide lines. Remove one of the glass plates, put the corresponding printout under the remaining glass plate, align with the gel and use as a guide to excise a gel slice from each lane, containing RNAs 30-55 nt long, or other desired range.
Further slice each piece of gel into 15-25 smaller pieces, transfer them to a 1.5 ml tube, and add 700 µl of TE buffer supplemented with 0.1% SDS. Mix on a rotator at 4°C for several hours or overnight. Centrifuge for 1 min at maximum speed, then carefully transfer the TE buffer into a fresh tube. We use 200-µl flat tips to avoid transferring small acrylamide pieces with the buffer. Add 7.5 µg of GlycoBlue, 75 µl of 3M sodium acetate pH 5.4, and mix. Add 750 µl of isopropanol, mix again, and precipitate overnight at -20 ˚C. Centrifuge for 30 min at 20,000 x g, 4 ˚C. Wash the pellets with 1 ml of 75% ethanol. Remove the ethanol completely, air dry the pellet for a maximum of 2 min. Dissolve in 6 µl DEPC treated MilliQ-$H_2O$ on ice.

**7. L3 linker ligation:**

Mix 6 µl dissolved RNA in 20-µl reaction also containing 1x RNA ligase1 buffer (New England BioLabs), 1 mM ATP (New England BioLabs), 16.9% PEG8000 (New England BioLabs), 15 U of T4 RNA ligase1 (New England BioLabs), 10 Ul RNaseOUT, and 75 ng (10.75 pmole) of L3 linker. Incubate at 16 °C for 16 hours. Perform extraction with acidic phenol as in step 2, and dissolve the dried pellet in 20 µl DEPC-treated MilliQ-water.

**8. Reverse transcription:**

Mix 3 µl of 10 mM dNTP mix (New England BioLabs) and 6 µl of 10 µM Rclip primer with an appropriate in-line barcode with 20 µl of the dissolved RNA. The primer is thus in over 5-fold excess of the L3 linker. Incubate in thermal cycler for 5 min at 70°C with lid heating. Take the tube from 70°C and place directly on ice for at least

1 min. Set the cycler at 25°C and equilibrate the samples at this temperature for 30-60 sec. Add a 31 µl mix containing 3 µl of 200 U/µl SuperScript IV Reverse Transcriptase (Thermo Fisher Scientific), 12 µl of the supplied 5x reaction buffer, 3 µl of 100 mM DTT, 3 µl of 40 U/µl RNaseOUT, and 10 µl of DEPC-treated MilliQ-water. Mix and incubate at 25°C for 5 min, then for 10 min at 42°C, and for another 15 min at 48°C. Transfer the reverse transcription reaction to a 1.5 ml tube, and perform deproteinization as in step 2, extracting with alkaline phenol:chloroform:isoamyl alcohol (125:24:1), pH 7.7–8.3 (Millipore-Sigma). Dissolve the pellet in 5 µl MilliQ-water

**9. Size selection of cDNA:**

Excise cDNAs from gel as in step 6. To select cDNAs of RNA-L3 linker templates, excise 86-111 nt (cDNA length = 56 nt + RNA fragment length). Dissolve cDNA pellets in 6.5 µl of MilliQ-water.

**10. Circularization of the cDNA:**

Add 1.5 µl of the following mix in a PCR tube: 0.8 µl of 10x Circligase II buffer, 0.4 µl of 50 mM $MnCl_2$, and 0.3 µl of 100 U/µl CircLigase II ssDNA ligase (LGC Biosearch Technologies). Add 6.5 µl of cDNA solution and mix by pipetting. Incubate in a thermal cycler at 60°C for 60 min with lid heating.

**18. Linearization at the BamHI site:**

Add 30 µl of the following mix: 4 µl of 10x FastDigest buffer (Thermo Fisher Scientific), 0.9 µl of 10 µM cut_oligo, and 25.1 µl of MilliQ-water. Incubate for 4 min at 95 ˚C, then decrease the temperature by 1°C every 60 sec until reaching 37°C. Add 2 µl of FastDigest BamHI (Thermo Fisher Scientific), mix and incubate for 30 min at 37°C.

Transfer the sample to 1.5 ml tube, adjust the volume to 200 µl, 5 mM EDTA, and perform phenol extraction with alkaline phenol:chloroform: isoamyl alcohol mix as in step 8. Dissolve the dried pellet in 22 µl of MilliQ-$H_2O$.

**19. Analytical PCR to determine the optimum number of cycles for preparative amplification:**

Prepare a 42-µl PCR reaction mix containing 2 µl of single stranded DNA template, 0.02U/ µl of Phusion Hot Start II high-fidelity DNA polymerase and 1x Phusion HF buffer (Thermo Fisher Scientific), 0.2 mM dNTP mix, and 0.2 µM of each P5solexa and P3solexa primers. Split this mix into four aliquots of 10 µl each in PCR tubes. Prepare a negative control the same way, adding water instead of template.

Amplify one aliquot from each PCR mix for 12, 16, 20, and 24 cycles using the following parameters: initial denaturation: 98°C for 30 sec, cycle (denaturation 98°C for 7 sec, annealing 64°C for 20 sec, extension 72°C for 15 sec), final extension for 5 min at 72°C, cool down to 4 ˚C.

Using a separate set of pipettes and a separate bench if available, run these samples on a 2% agarose gel containing 0.5x TBE and 0.5 µg/ml Ethidium bromide. For each sample, calculate the number of cycles that will produce about 50-200 ng of PCR product from the rest of the template.

**20. Preparative PCR:**

Prepare 30 µl of PCR reaction mix the same way as above, using 20 µl of single stranded DNA template. Amplify by PCR using the parameters described above and the number of cycles determined at the analytical step. Run the PCR reactions on a 2% low-melting point agarose gel and excise gel bands containing products ranging from 160 to 190 bp (PCR product length = 132 bp + RNA fragment length). Extract DNA from the agarose gel slice using a Zymoclean Gel DNA recovery kit (Zymo Research). Elute DNA from the column with 10 µl of EB buffer (Qiagen).

Determine the concentration of DNA by Qubit using the dsDNA BR Assay Kit (Thermo Fisher Scientific). Prepare 5-20 µl of sequencing library, containing 10 nM DNA and 0.1% Tween-20 in buffer EB. Multiple PCR products can be mixed together if they were prepared with RT primers bearing different barcodes.

Sequence in Illumina high-throughput sequencer at single end 100 nt.
